# Organization of cortical and thalamic input to inhibitory neurons in mouse motor cortex

**DOI:** 10.1101/2021.07.08.451716

**Authors:** Sandra U. Okoro, Roman U. Goz, Brigdet W. Njeri, Madhumita Harish, Catherine F. Ruff, Sarah E. Ross, Charles R. Gerfen, Bryan M. Hooks

**Author notes:** Correspondence: Dr. Bryan M. Hooks, 200 Lothrop Street BSTWR Suite W1458 Pittsburgh, PA 15213.

## Abstract

Intracortical inhibition in motor cortex (M1) regulates movement and motor learning. If inhibitory cell types and cortical laminae targeted by cortical and thalamic afferents differ, then these afferents play different roles in regulating M1 output. We quantified input to two classes of M1 interneurons, parvalbumin+ (PV) fast-spiking cells and somatostatin+ (SOM) low-threshold-spiking cells, using monosynaptic rabies tracing. We then compared anatomical connectivity and functional connectivity based on synaptic strength from sensory cortex and thalamus. Functionally, each input innervated M1 interneurons with a unique laminar profile. Different interneuron types were excited in a distinct, complementary manner, suggesting feedforward inhibition proceeds selectively via distinct circuits. Specifically, somatosensory cortex (S1) inputs primarily targeted PV+ neurons in upper layers (L2/3) but SOM+ neurons in middle layers (L5). Somatosensory thalamus (PO) inputs targeted PV+ neurons in middle layers (L5). In contrast to sensory cortical areas, thalamic input to SOM+ neurons was equivalent to PV+ neurons. Thus, long-range excitatory inputs target inhibitory neurons in an area and cell type-specific manner which contrasts with input to neighboring pyramidal cells. In contrast to feedforward inhibition providing generic inhibitory tone in cortex, circuits are selectively organized to recruit inhibition matched to incoming excitatory circuits.

## INTRODUCTION

Motor cortex (M1) integrates long-range input from sensory cortex (S1) and thalamus to plan and control movement. Excitatory connectivity to excitatory neurons has been studied, but how different interneuron types are recruited by long-range input has received less attention. Cortical and thalamic sensory inputs arrive in M1, resulting in sensory activity (Ferezou et al., 2007; Hatsopoulos and Suminski, 2011; Huber et al., 2012; Murray and Keller, 2011), consistent with a role for M1 in active sensation (Hill et al., 2011) and sensorimotor learning (Asanuma, 1981). Input from S1 (Hoffer et al., 2003) excites pyramidal neurons primarily in upper layers of M1 (Kaneko et al., 1994a; Mao et al., 2011). Somatosensory input to M1 also originates from higher-order sensory thalamus, including the posterior nucleus (PO) (Deschenes et al., 1998; Harris et al., 2019; Ohno et al., 2012). These two classes of input (S1 and PO) both most strongly excited pyramidal neurons of L2/3 and L5A (Hooks, 2017; Hooks et al., 2013; Mao *et al*., 2011). L2/3 and L5A neurons then provide descending input to pyramidal neurons in L5B that regulate movement (Anderson et al., 2010; Hooks et al., 2011; Kaneko et al., 1994b; Kiritani et al., 2012; Weiler et al., 2008).

A diverse range of cortical GABAergic interneuron types exist, playing an important role in regulating cortical excitation (Tremblay et al., 2016). Three major classes include parvalbumin-expressing (PV+), somatostatin-expressing (SOM+), and 5-HT3a receptor-expressing (5HT3aR+) interneurons (Lee et al., 2010). Of these, PV+ and SOM+ cells are found across the width of M1 from L2 to L6 (Fig. 1) (Lee *et al*., 2010), while 5-HT3aR+ cells are restricted to the upper layers of cortex. PV+ interneurons include fast spiking interneurons, which target perisomatic regions of pyramidal neurons, and SOM+ interneurons, which include low threshold spiking interneurons (Cauli et al., 1997; Gibson et al., 1999; Kawaguchi and Kubota, 1997; Kubota, 2014). A subset of SOM+ interneurons, Martinotti cells, targets apical dendrites of pyramidal cells (Wang et al., 2004). Differences in subcellular targeting of excitatory cells contributes to different roles in regulating cortical output. Thus, understanding how different classes of interneuron are activated by distinct excitatory inputs is an important step in understanding how feedforward inhibition is recruited by different cortical afferents.

**Figure 1.**
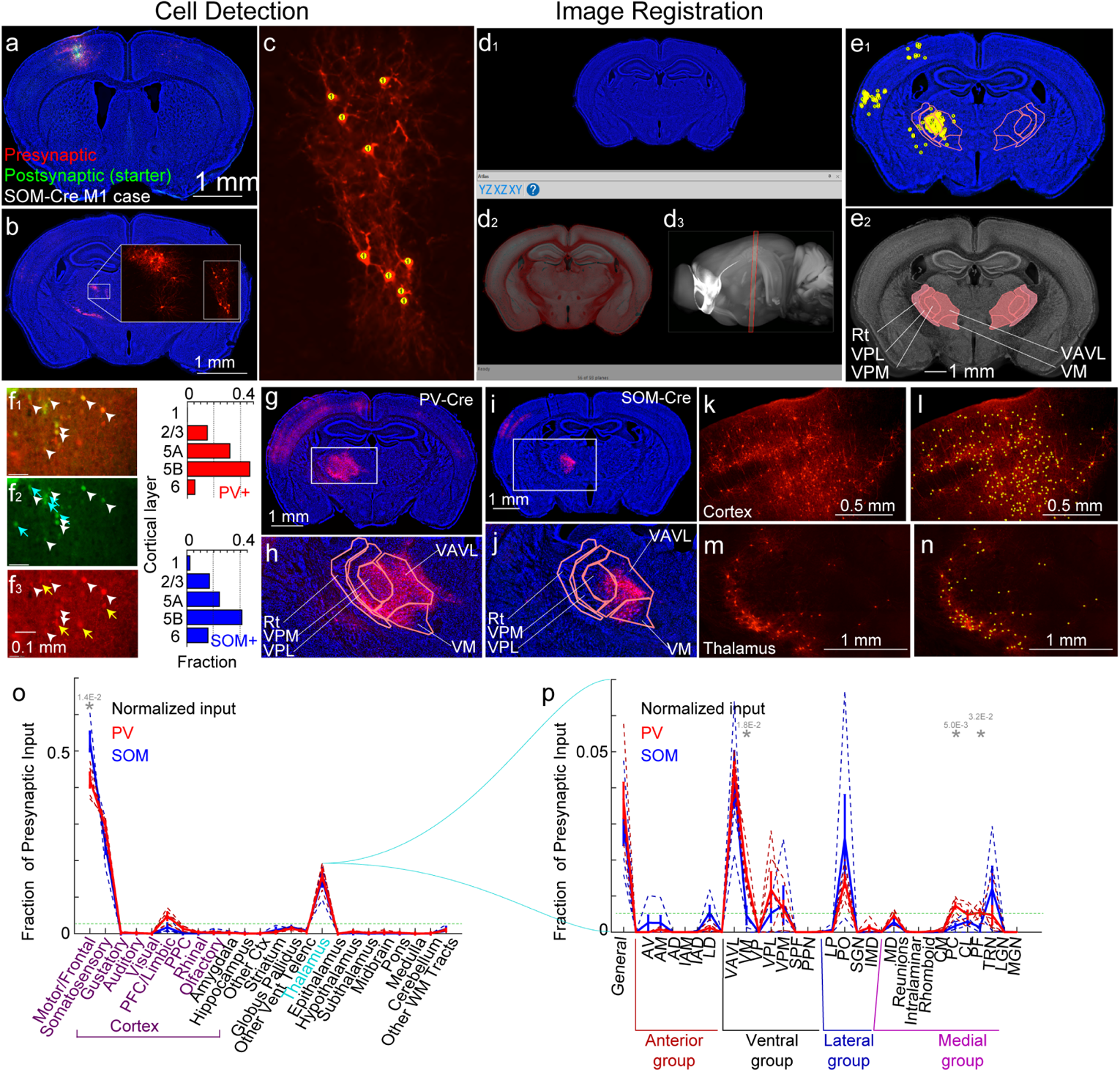
Monosynaptic Retrograde Tracing of Inputs to Motor Cortex PV+ and SOM+ Neurons. (a) Injection site in M1 of a SOM-Cre mouse injected with AAV-DIO-TVA-RG-EGFP (green) and EnvA pseudotyped N2c RABV-tdTomato (red). Neurotrace Blue labels the tissue structure. (b) tdTomato-labeled thalamic neurons (inset). (c) tdTomato+ (red) somata detected in NeuroInfo. Markers are yellow. (d) Section registration in NeuroInfo. (d1) NeuroTrace Blue used as a structural marker in imaged sections. (d2-d3) Current reference section (teal, d2) and brain (black and white, d3) showing the reference atlas. The reference section (d2) can be aligned to the current section (d1; overlaid in red in d2) by adjusting the scale, or shifting and rotating the plane in which the reference brain is sectioned (control panel in e). The current plane orientation for the reference brain is shown as the red box in d3 (slightly offset from coronal). A nonlinear transform for each section is used to register each object in the current image to the reference brain coordinates. Detected neurons (from b-c) are then transformed to the reference atlas coordinate system. (e) Coronal sections showing the Allen CCF 3.0 and locations of S1 and motor and somatosensory thalamic nuclei (Rt, thalamic reticular nuc.; VAVL, ventroanterior/ventrolateral nuc.; VM, ventromedial nuc.; VPL, ventroposterolateral nuc.; VPM, ventroposteromedial nuc.). The positions of retrogradely labeled presynaptic neurons are marked with a yellow dot (top image) in an example SOM-Cre mouse injected in M1. Presynaptic neurons within ±0.1 mm of the plane are shown. The bottom image (e2) is labeled to indicate registered thalamic areas. (f) Injection site with starter cells in a SOM-Cre mouse. (f1) Merged image of green (f2) and red (f3) channels, showing yellow starter cells (white arrowheads in f1-f3) and singly labeled cells (blue arrows in f1 and yellow arrows in f3). (f2) Green, TVA+ neurons (putative starter cells); (f3) Red, tdTomato+ neurons (putative presynaptic neurons). Right, laminar distribution of PV+ and SOM+ somata, quantified as fraction of the total by layer. (g-h) Example section (g, expanded in h) showing retrograde label in S1 and thalamus from M1 starter cells in a PV-Cre mouse. Note S1 label and thalamic label. (i-j) Example section showing retrograde label in S1 and thalamus from M1 starter cells in a SOM-Cre mouse. Note S1 label and thalamic label. (k-n) Cell detection examples in a SOM-Cre mouse. In cortex (k), detected cell somata are labeled with yellow circles (l). Example images for thalamus (m-n). (o) Quantification of presynaptic neuron location from brains aligned to the Allen CCF V3.0. Fraction of the total presynaptic neuron population (±SEM) plotted for all M1 injections in PV-Cre (N=5, red) and SOM-Cre (N=4, blue) mice. Dashed lines represent individual cases; solid line represents mean. Green dashed line, threshold for testing. *, t-test p<0.05 (gray *, not significant when controlling for false discovery rate) (p) Quantification of thalamic inputs for all M1 injections in PV-Cre (N=5, red) and SOM-Cre (N=4, blue) mice. Data plotted as in (o), subdivided into distinct thalamic nuclei.

We tested whether the connectivity of long-range inputs to PV+ and SOM+ interneurons in M1 follows the pattern from sensory cortex studies. Because ascending thalamic inputs in sensory cortical areas principally target PV+ interneurons, we hypothesized that M1 PV+ interneurons would similarly be the major recipients of thalamic input. These conclusions are derived from studies of the connectivity of principal thalamic nuclei to cortical layer 4 (Cruikshank et al., 2007; Cruikshank et al., 2010), which is absent in M1. First, we quantified the overall pattern of input to specific interneuron types using monosynaptic rabies tracing. Then, because this method is not layer specific, we measured monosynaptic excitation to M1 interneurons functionally using channelrhodopsin-2 (ChR2)-assisted circuit mapping (Cruikshank *et al*., 2010; Petreanu et al., 2007; Petreanu et al., 2009) to assess synaptic input strength as a function of laminar depth and specific interneuron type across layers in M1. In contrast to the strong preference of thalamocortical input for PV+ interneurons in layer 4 of sensory cortex (Cruikshank *et al*., 2007; Cruikshank *et al*., 2010), both methods found thalamic input to SOM+ neurons that was nearly as strong as PV+ neuron input. The laminar pattern, however, differed. Corticocortical afferents from S1 also targeted interneurons differently across cell types: among PV+ neurons, L2/3 cells were most strongly excited, while for SOM+ neurons, input arrived strongly in middle layers (L5). Furthermore, comparing inputs from S1 and thalamus, PV+ neurons were targeted in a complementary fashion, with S1 exciting upper layers (L2/3) and thalamus targeting middle layers (L5). Because S1 and PO targeted excitatory neurons in similar layers (Hooks et al., 2015; Hooks *et al*., 2013), the complementary pattern of input to PV+ neurons suggests that feedforward inhibition does not simply silence cortex, but integrates into local circuits in a specific fashion.

## RESULTS

### Anatomical tracing of long-range inputs to PV+ and SOM+ neurons

Incoming cortical and thalamic excitation to M1 directly excites pyramidal neurons (Hooks *et al*., 2013; Mao *et al*., 2011) and interneurons, evoking disynaptic feedforward inhibition. The interneurons carrying this feedforward inhibition likely include PV+ basket cells as well as SOM+ neurons. Conventional retrograde tracers can identify presynaptic cortical (Dum and Strick, 2002; Hoffer *et al*., 2003; Reep et al., 1990; Rouiller et al., 1993) and thalamic inputs (Deschenes *et al*., 1998; Kuramoto et al., 2009; Kuramoto et al., 2015; Ohno *et al*., 2012) to M1 (Hooks *et al*., 2013; Oh et al., 2014; Zingg et al., 2014), but do not differentiate between cell types in M1 receiving the input. To quantify how specific populations of interneurons are targeted by incoming cortical and thalamic afferents, we used genetically identified interneurons as starter cells for monosynaptic retrograde tracing with rabies virus (Callaway and Luo, 2015; Wickersham et al., 2007b). This enabled us to quantitatively compare presynaptic labeling across starter neuron types (PV+ and SOM+ neurons) as well as across cortical areas.

Using PV-Cre or SOM-Cre mice, we infected Cre+ interneurons with AAV-DIO-TVA-EGFP-B19G to label potential starter neurons and make them express both TVA receptor (to become infected with EnvA pseudotyped rabies virus) as well as the rabies coat protein B19G (Fig. 1f). The starter cells in these injections spanned cortical laminae (L2-6). After two weeks to express the receptor, EnvA-pseudotyped CVS-N2c G-deleted rabies expressing tdTomato (Reardon et al., 2016) was injected in forelimb motor cortex (fM1, N=5 for PV+, N=4 for SOM+ mice; see Table 1). Fixed brains were sectioned, immunoamplified, and imaged. The whole brain image stacks were reconstructed (Fig. 1a). Cell detection was then performed for tdTomato+ neurons in NeuroInfo (MBF Bioscience) in each plane using an artificial neural network (Fig. 1b-c; Supp. Fig. 1). The image stack was aligned to the Allen CCF V3.0 (Kuan et al., 2015; Oh *et al*., 2014) using the structural label Neurotrace (blue, Fig. 1d-e), enabling the assignment of voxels containing structures and cells to defined cortical and thalamic regions (Fig. 1g-n; see Supp. Fig. 2 for detail of alignment in thalamus). fM1 was used to compare the data to results from the Brain Initiative Cell Census Network (BICCN) studies (Muñoz-Castaneda, 2020). This approach was limited since injections did not restrict starter PV+ or SOM+ neurons to a single cortical layer.

**Table 1.** Injection coordinates. Anterior/posterior (A-P) axis reported relative to bregma (positive values anterior to bregma). Medial/lateral (M-L) axis reported relative to the midline. Dorsal/ventral (D-V) axis, depth from pia. Injections were made at both depths in cortex. Distances in mm.

| Target | A-P | M-L | D-V |
| --- | --- | --- | --- |
| Primary whisker motor cortex (vM1) | +1.1 | 0.9 | 0.5/0.8 |
| Posterior thalamus (PO) | -1.2 | 1.3 | 2.75 |
| Primary whisker somatosensory cortex (vS1) | -0.6 | 3.0 | 0.5/0.8 |
| **For rabies tracing** |  |  |  |
| Secondary motor cortex (M2) | +2.6 | 1.0 | 0.5/0.8 |
| Forelimb motor cortex (fM1) | +0.6 | 1.5 | 0.5/0.8 |
| Primary whisker somatosensory cortex (vS1) | -0.6 | 3.0 | 0.5/0.8 |

The distribution of presynaptic neurons was quantified by normalizing the number of cells detected in each brain region as a fraction of the total number of cells detected (Fig. 1o-p). The distributions were repeatable across different cases in the same brain region (Fig. 1o-p and Supp. Fig. 3), which indicated that the NeuroInfo alignment was effective across individual brains and that the retrograde tracing method was reliable. To minimize excessive comparisons, we limited comparisons to brain regions where at least one area exceeded a detection threshold (dotted line in Fig. 1o-p). Overall, the pattern was quite similar for inputs to PV+ and SOM+ neurons in M1. Most presynaptic inputs were found in cortex (Fig. 1o), in particular frontal and motor areas on the dorsal surface of cortex, as well as somatosensory cortex, which is strongly reciprocally connected to mouse M1 (Hooks, 2017; Mao *et al*., 2011). About 20% of inputs for both interneuron classes were localized to thalamus, consistent with similar corticocortical and thalamocortical input to these cell types. We further examined the thalamocortical data by subdividing labeled neurons by thalamic nuclei (Fig. 1p). Although the data showed that the thalamic nuclei associated with M1 based on prior anatomic studies (Deschenes *et al*., 1998; Kuramoto *et al*., 2009; Kuramoto *et al*., 2015; Ohno *et al*., 2012) were robustly represented, including VAVL, VM, and PO, we did not detect significant differences between number of neurons projecting to PV+ or SOM+ neurons. The ability to detect such differences was limited by correction for false detection rate (Benjamini et al., 2009), with non-significant trends in VM and two medial nuclei (PC and PF).

We extended the retrograde viral tracing study to two other cortical areas strongly reciprocally connected with motor areas, primary somatosensory cortex and frontal cortex (M2). Experimental procedures were identical, except for the targeting of viral injections to different cortical areas in PV-Cre and SOM-Cre mouse lines. This gave a richer dataset for comparison. We first made comparisons across cortical areas of starter cell within a given interneuron line (Fig. 2b-d). For SOM+ neurons, the major areas labeled were motor cortex and frontal cortex (greater in fM1 and M2 starter cases than in vS1 starter cases) and somatosensory cortex (greater in vS1 starter cases), with a relatively uniform fraction of thalamic input (~20%) across neocortical areas). PV+ input was similar, with the additional note of significantly greater input from limbic areas (mostly orbital cortex) to M2 than to other areas. Comparison of input from specific thalamic nuclei was considerably different across injection sites. In general, the pattern of thalamic label could be used to infer the cortical target, with a reasonably large number of nuclei providing a specific pattern of cortical input. For example, in cases using PV+ starter neurons, vS1 injections could be identified by more robust VPM and PO labeling. fM1 injections had strong input from VAVL and VM as well as PO and midline nuclei. M2 input was similar to fM1 in receiving strong VAVL and VM input, but also received significant MD input (which vS1 and fM1 did not) as well as input from midline nuclei including PC and PF. This is consistent with substantial differences in thalamocortical targeting across cortical areas. However, in comparing input to different cell types within a given cortical area, few significant differences were detected. For M2, PV+ interneurons labeled more thalamic cells in VM and PC compared to SOM+ neurons in M2. In vS1, SOM+ interneurons labeled more VAVL neurons than PV+ neurons. For most nuclei, there were no detectable differences. In general, this was consistent with much larger differences in thalamic label for cortical areas than for specific cell types within a cortical area.

**Figure 2.**
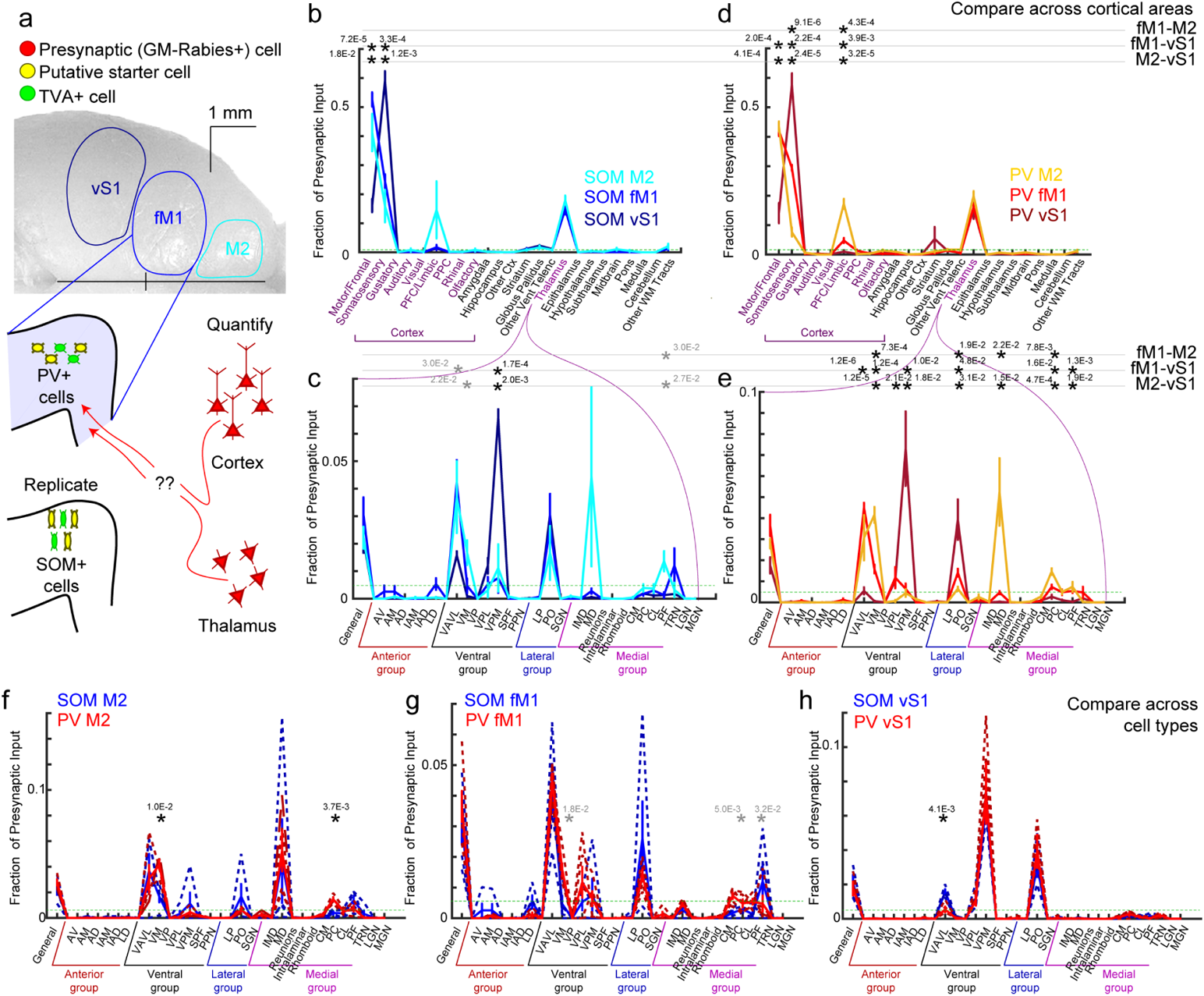
Monosynaptic Retrograde Tracing of Inputs to PV+ and SOM+ Neurons Across Cortical Areas. (a) Schematic showing three dorsal cortical areas used for retrograde tracing from PV+ and SOM+ interneurons. In PV-Cre or SOM-Cre mice, injection of AAV expressing DIO-TVA-EGFP-B19G labels potential starter cells green in a Cre-dependent manner. This vector also expressed the TVA receptor to permit rabies uptake. Subsequent injection of EnvA-pseudotyped CVS-N2c-tdTomato rabies labels starter cells yellow (red+green), at left in cartoon and labels potential presynaptic partners red (at right in many cortical and thalamic areas). (b-c) Quantification of presynaptic neuron location for injections in SOM-Cre (N=4, M2; N=4, fM1; N=4, vS1; cool colors) mice. Whole brain image stacks were aligned to the Allen CCF V3.0. Presynaptic neuronal somata were detected and coordinates in CCF assigned to brain regions. Mean fraction of the total presynaptic neuron population (±SEM) plotted as in Fig. 1. Quantification of inputs to all brain regions (b) and to thalamic nuclei (c). (d-e) Data from injections in PV-Cre (N=4, M2; N=5, fM1; N=5, vS1; hot colors) plotted as in (b-c). (f-h) Data of (b-e) replotted to compare input to SOM+ (blue) and PV+ (red) neurons in the same cortical locations: M2 (f), fM1 (g), and vS1 (h). Dotted lines represent individual cases; solid lines represent mean. Green dashed horizontal lines in b-h represent threshold for statistical comparison. Black * represent significant differences (p value as indicated) after correcting for FDR. Gray * represent comparisons not significant after correction for FDR.

### Synaptic mapping of long-range excitatory inputs from S1 and thalamus to M1

M1 directly excites pyramidal neurons (Hooks *et al*., 2013; Mao *et al*., 2011). Our data showed that these inputs were capable of evoking disynaptic feedforward inhibition by recording from pyramidal neurons in M1 in brain slice while exciting ChR2+ axons from S1 at −70 mV and +0 mV and quantifying the EPSC and IPSC amplitude, respectively (Fig. 3). The feedforward inhibition at +0 mV onset ~2 ms slower than the excitation (Fig. 3a). Across M1 layers, excitation is matched to inhibition, with similar ratios across layers. The interneurons carrying this feedforward inhibition are unknown, but are presumed to include PV+ basket cells. Further, it is likely that the GABAergic neurons carrying the disynaptic inhibition reside mainly in the same layer as the pyramidal neuron soma (Katzel et al., 2011). To identify the source of this disynaptic inhibition, we then quantified how specific populations of interneurons are targeted by incoming cortical and thalamic afferents. This experiment will also permit a direct comparison of functional synaptic connectivity by synaptic circuit mapping and anatomical tracing by retrograde viruses. We labeled axons from S1 or PO with ChR2-mVenus. These axons arborized in distinct layers of cortex (Fig. 3c-d). We selected two populations of Cre-driver mice (PV-Cre and SOM-Cre) crossed to a tdTomato reporter (Ai14) (Madisen et al., 2010) in order to label genetically defined interneurons across the thickness of M1. We then targeted whole-cell recordings to tdTomato+ interneurons in M1 brain slices in the field of mVenus+ axon terminals. We used a 470 nm light to excite ChR2+ axon terminals while recording in TTX, CPP, and 4-AP (sCRACM) (Petreanu *et al*., 2007; Petreanu *et al*., 2009). TTX prohibited action potentials, ensuring inputs measured were monosynaptic. 4-AP slowed axon repolarization, allowing the ChR2-evoked EPSC to occur (Fig. 3g-h). Because we were concerned that axons targeting different cell types might vary in their responsiveness to 4-AP, we confirmed that addition of 4-AP in the presence of TTX restored synaptic release with similar effectiveness. In mapping experiments, a single interneuron was recorded while stimulating the brain slice with a 470 nm laser. A 12×26 grid (Fig 3j) was used for stimulation, resulting in evoked EPSCs (Fig. 3l) that differed in amplitude depending on the location of the laser stimulus. A heat map showing the size of the response was then plotted on top of the cell location in the slice (Fig. 3k). We compared each pair of interneurons in different layers recorded in the same slice, summing the EPSC amplitude across the map. This measure of input strength was used to compare EPSC amplitude across layers (Fig. 3m). Thus, our comparisons of input strength (Fig. 4–7) required pairwise comparison of neurons recorded in the same slice to normalize for ChR2 expression between animals, and nonparametric statistics were used to avoid assumptions about the distribution of connection strength.

**Figure 3.**
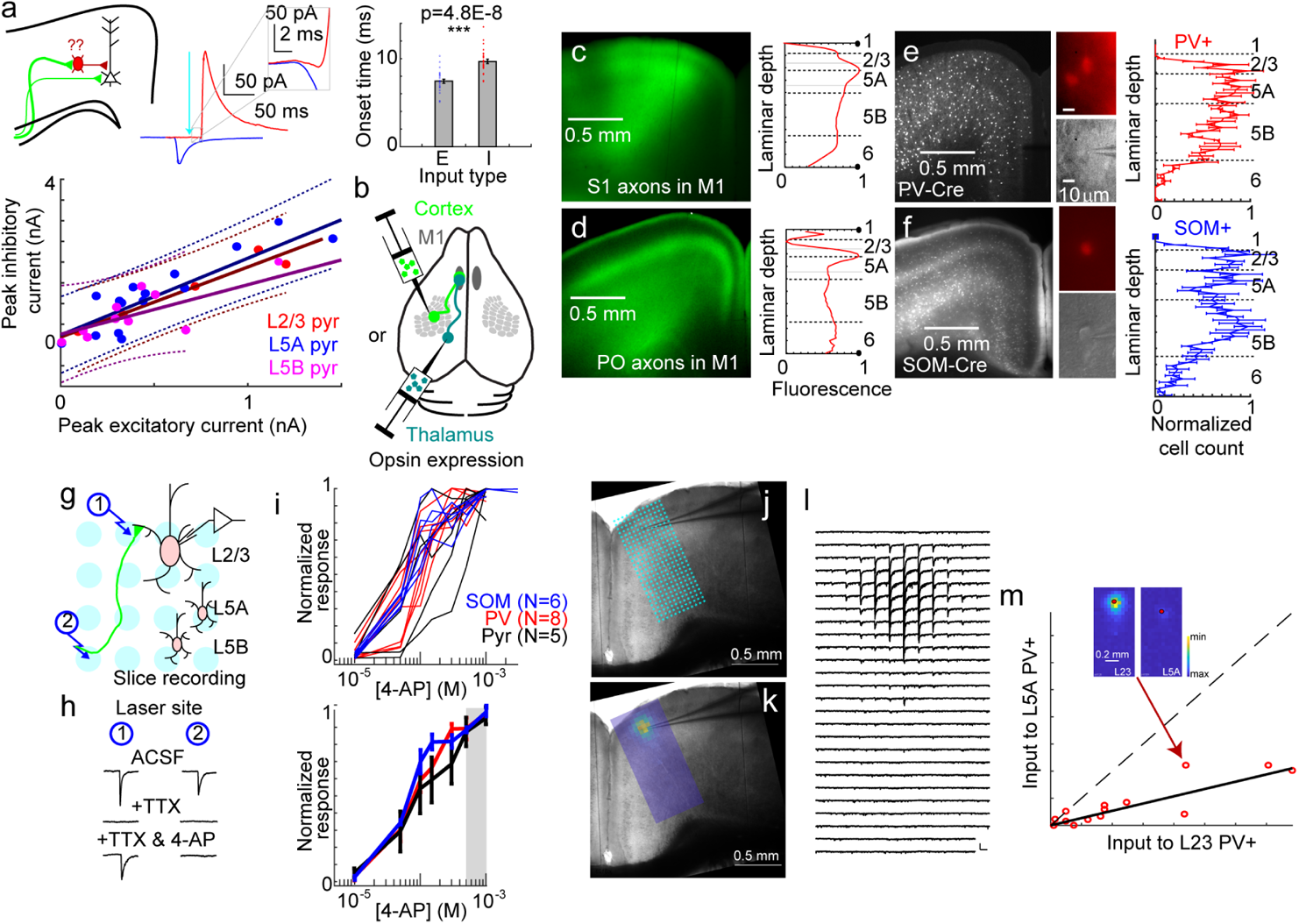
Mapping long-range excitatory inputs to mouse motor cortex interneurons. (a) Interneurons carry feedforward inhibition in M1. Whole-cell recordings at −70 mV (blue) and 0mV (red) show optically induced feedforward inhibition from S1 inputs to M1 (bottom left) with a short latency difference (inset). Difference in latency between EPSC (blue) and IPSC (red) is plotted to compare onset (mean and SE; N=29). EPSC and IPSC amplitude are roughly proportional (right) across all cortical layers (bottom right). Linear fit and 95% confidence interval (dashed lines) plotted. (b) AAV vectors expressing optogenetic activators (ChR2 or Chronos) were stereotaxically injected into cortex (S1, light green) or thalamus (PO, dark green). (c) mVenus+ axons from S1 arborize across layers of M1 in a columnar projection (left, 4x image). Fluorescence is highest in L5A (right, brightness quantified across cortical layers). (d) mVenus+ axons from PO arborize across layers of M1 in two main layers (left). Fluorescence is highest in L1 and L5A (right). (e) PV-Cre mice were crossed to a tdTomato reporter (Ai14), labeling interneurons across M1 in L2-6. Postsynaptic interneurons were recorded across all layers for comparison. Inset (60x, fluorescence image, top, and corresponding brightfield image, bottom) shows patch pipette on tdTomato+ neuron in PV-Cre x Ai14 mouse. Normalized laminar distribution of PV+ somata (N=5 slices) measured at right. (f) SOM-Cre mice were crossed to a tdTomato reporter (Ai14), labeling interneurons across M1 in L2-6. L1 signal represents tdTomato+ axons from SOM+ neurons. Postsynaptic interneurons were recorded across all layers for comparison. Normalized laminar distribution of SOM+ somata (N=5 slices) measured at right. (g) Targeted whole cell recordings in brainslice were made from interneurons while stimulating with a 470nm laser at different points, shown in light blue. Full grid illustrated as in (j). (h) 470nm stimulation excites axons at example points 1 & 2, but TTX addition extinguishes responses. Addition of 4-AP permits local release at points where opsin+ axons contact interneuron dendrites (Point 1) but not distant points along the axon (Point 2). (i) Similar concentrations of 4-AP are required to restore synaptic responses regardless of postsynaptic neuron type (Pyr = pyramidal neuron). Individual cells plotted at top; average and SE plotted at bottom. Grey box represents [4-AP] used for mapping. (j) Typical input mapping experiments sampled a 12×26 point grid with points spaced at 50 mm and aligned to the pia. Brightfield image (4x) shows recording pipette in M1. (k) Heatmap shows stronger responses (red) near the soma of the recorded neuron. (l) Example traces for a L2/3 PV+ neuron show location of input. (m) Inset shows two maps to compare input between L2/3 and L5A PV+ interneurons. The summed synaptic input across the map is compared for each cell pair recorded in the same slice. Arrow indicates a point from one L2/3-L5A pair. Dashed line represents y=x (similar input to both layers); solid line represents the geometric mean of the L2/3 and L5A input strength, presented as a line of equivalent slope.

**Figure 4.**
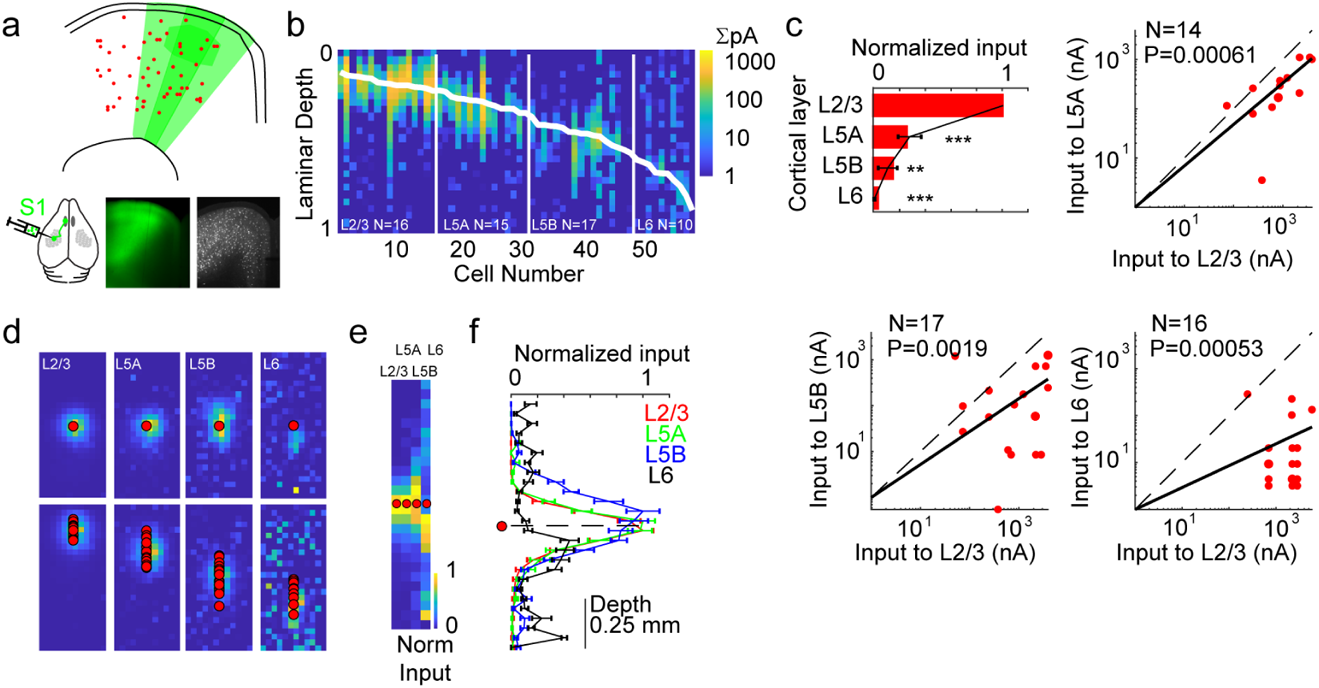
S1 input to M1 targets upper layer PV+ neurons. (a) Cartoon showing PV+ neurons (red) in GFP+ axons from S1. (b) S1 input to M1 PV+ neurons across layers (N=58 total). Each cell is represented as a single column (vector), with the rows of the input mapped summed and aligned to the pia. Diagonal white line represents the soma location for that cell. Vertical white lines represent layer divisions. Synaptic strength (summed, in nA) for each location in depth are represented by the heatmap (scale at right). Laminar depth is normalized to 0=pia, 1=white matter. (c) Strength of synaptic input. The bar represents the geometric means of the amplitude ratio, normalized to the layer receiving the strongest input (L2/3). The overlaid graph shows the mean ratio and SD (based on 10000 replicate bootstrap). Adjacent to the summary, three graphs for comparison of input strength across neurons in different cortical layers. Each point represents input to a pair of neurons in the same slice (circle for each neuron). Dashed line represents unity. N, Number of pairs; p value, Wilcoxon signed rank test. (d) Maps of synaptic input location in the dendritic arbor for PV+ neurons in each layer. Top row, normalized soma-centered map (maps registered to soma center across cells). Bottom row, normalized pia-aligned maps. Normalized maps are noisy when input is weak. (e,f) Input location summarized for all four layers. Normalized mean input maps were averaged into a vector (e) and aligned to the soma (red circle). These were graphed with mean and SD (f), showing input relative to the soma in 50 mm bins. Dashed line indicates soma depth. *=p<0.05, **=p<0.01, and ***=p<0.001.

**Figure 5.**
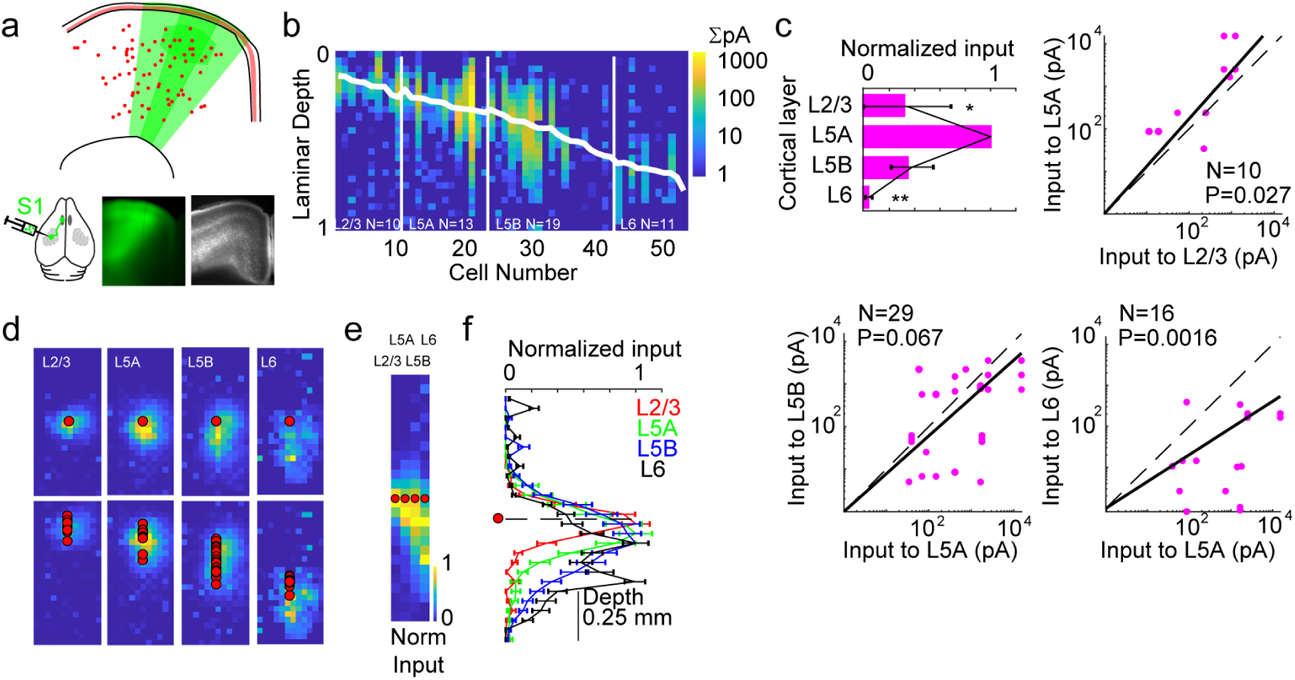
S1 input to M1 targets middle layer SOM+ neurons. (a) Cartoon showing SOM+ neurons (red) in GFP+ axons from S1. (b) S1 input to SOM+ neurons across layers (N=53 total). Each cell is represented as a single column (vector), with the rows of the input mapped summed and aligned to the pia. Diagonal white line represents the soma location for that cell. Vertical white lines represent layer divisions. Synaptic strength (summed, in nA) for each location in depth are represented by the heatmap (scale at right). Laminar depth is normalized to 0=pia, 1=white matter. (c) Strength of synaptic input. The bar represents the geometric means of the amplitude ratio, normalized to the layer receiving the strongest input (L5A). The overlaid graph shows the mean ratio and SD (based on 10000 replicate bootstrap). Adjacent to the summary, three graphs for comparison of input strength across neurons in different cortical layers. Each point represents input to a pair of neurons in the same slice (circle for each neuron). Dashed line represents unity. N, Number of pairs; p value, Wilcoxon signed rank test. (d) Maps of synaptic input location in the dendritic arbor for SOM+ neurons in each layer. Top row, normalized soma-centered map (maps registered to soma center across cells). Bottom row, normalized pia-aligned maps. Normalized maps are noisy when input is weak. (e,f) Input location summarized for all four layers. Normalized mean input maps were averaged into a vector (e) and aligned to the soma (red circle). These were graphed with mean and SD (f), showing input relative to the soma in 50 mm bins. Dashed line indicates soma depth.

**Figure 6.**
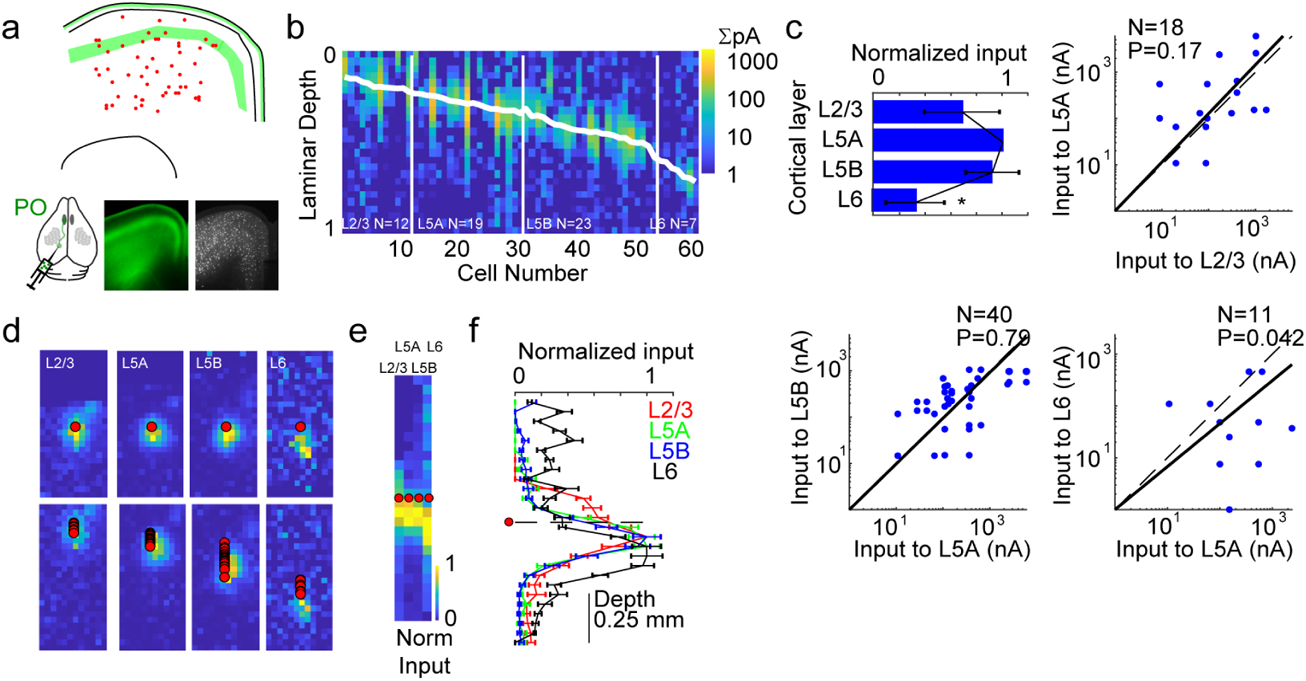
PO input to M1 PV+ neurons targets upper and middle layers. (a) Cartoon showing PV+ neurons (red) in GFP+ axons from PO. (b) PO input to PV+ neurons across layers (N=61 total). Each cell is represented as a single column (vector), with the rows of the input mapped summed and aligned to the pia. Diagonal white line represents the soma location for that cell. Vertical white lines represent layer divisions. Synaptic strength (summed, in nA) for each location in depth are represented by the heatmap (scale at right). Laminar depth is normalized to 0=pia, 1=white matter. (c) Strength of synaptic input. The bar represents the geometric means of the amplitude ratio, normalized to the layer receiving the strongest input (L5A). The overlaid graph shows the mean ratio and SD (based on 10000 replicate bootstrap). Adjacent to the summary, three graphs for comparison of input strength across neurons in different cortical layers. Each point represents input to a pair of neurons in the same slice (circle for each neuron). Dashed line represents unity. N, Number of pairs; p value, Wilcoxon signed rank test. (d) Maps of synaptic input location in the dendritic arbor for PV+ neurons in each layer. Top row, normalized soma-centered map (maps registered to soma center across cells). Bottom row, normalized pia-aligned maps. Normalized maps are noisy when input is weak. (e,f) Input location summarized for all four layers. Normalized mean input maps were averaged into a vector (e) and aligned to the soma (red circle). These were graphed with mean and SD (f), showing input relative to the soma in 50 mm bins. Dashed line indicates soma depth.

**Figure 7.**
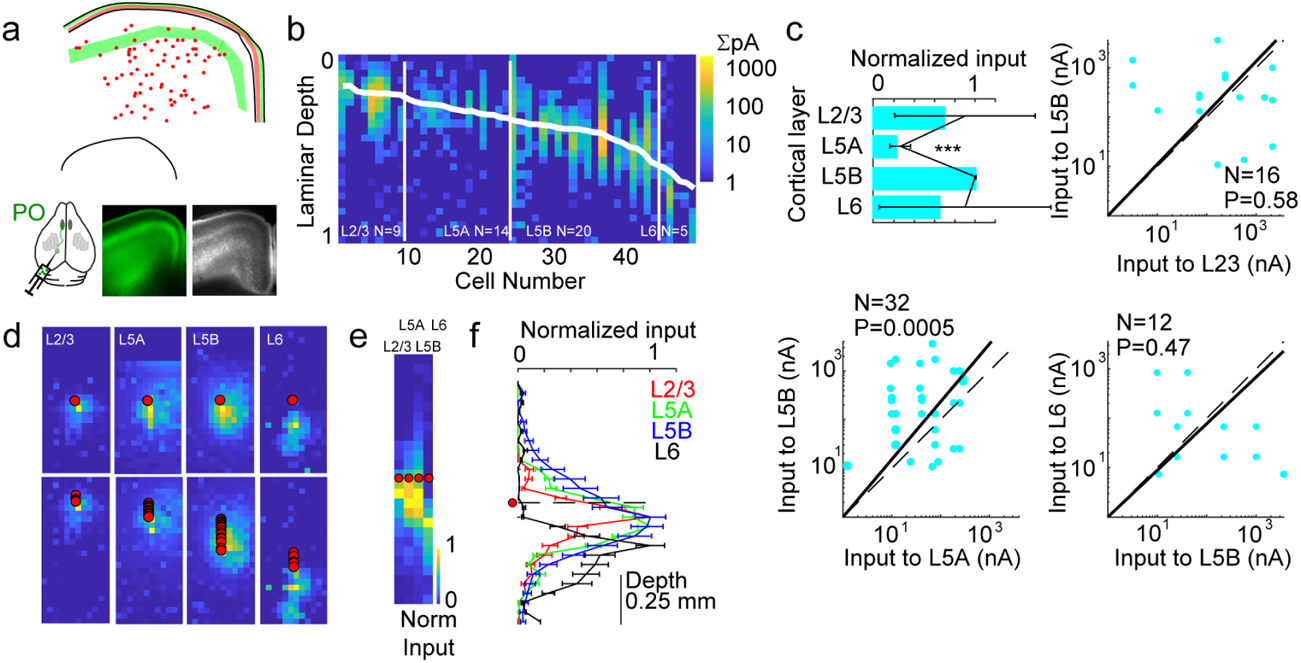
PO input to M1 SOM+ neurons is variable. (a) Cartoon showing SOM+ neurons (red) in GFP+ axons from PO. (b) PO input to SOM+ neurons across layers (N=48 total). Each cell is represented as a single column (vector), with the rows of the input mapped summed and aligned to the pia. Diagonal white line represents the soma location for that cell. Vertical white lines represent layer divisions. Synaptic strength (summed, in nA) for each location in depth are represented by the heatmap (scale at right). Laminar depth is normalized to 0=pia, 1=white matter. (c) Strength of synaptic input. The bar represents the geometric means of the amplitude ratio, normalized to the layer receiving the strongest input (L5B). The overlaid graph shows the mean ratio and SD (based on 10000 replicate bootstrap). Adjacent to the summary, three graphs for comparison of input strength across neurons in different cortical layers. Each point represents input to a pair of neurons in the same slice (circle for each neuron). Dashed line represents unity. N, Number of pairs; p value, Wilcoxon signed rank test. Plots show substantial heterogeneity. (d) Maps of synaptic input location in the dendritic arbor for SOM+ neurons in each layer. Top row, normalized soma-centered map (maps registered to soma center across cells). Bottom row, normalized pia-aligned maps. Normalized maps are noisy when input is weak. (e,f) Input location summarized for all four layers. Normalized mean input maps were averaged into a vector (e) and aligned to the soma (red circle). These were graphed with mean and SD (f), showing input relative to the soma in 50 mm bins. Dashed line indicates soma depth.

### S1 targeting of PV+ and SOM+ interneurons in M1

First, we recorded from tdTomato+ PV+ interneurons in the whisker region of M1 in PV-Cre x Ai14 mice. S1 inputs to M1 mainly targeted PV+ interneurons in L2/3. L2/3 PV+ neurons received more than twice as much input than any other layer. All PV+ neurons (N=58) in all layers were plotted for comparison (Fig. 4b), with each map reduced to a column vector by summing the input map across rows. Most responsive cells and the strongest responding cells were found in L2/3. A pairwise comparison of L2/3 with other PV+ neurons recorded in the same slice showed that L5A neurons received only about 30% the input of L2/3 PV+ neurons (P<0.001). Similarly, input to L5B and L6 was even weaker (Fig. 4c). This input was mostly centered around the interneuron soma, as shown in maps where the grid is aligned to the soma (Fig. 4d-e). For layers with weaker input, noise is increased in normalized input maps (for example, L6 in Fig. 4d).

We repeated this experiment for S1 inputs to SOM+ interneurons, using SOM-Cre x Ai14 mice (Fig. 5). As in PV-Cre mice, SOM-Cre labeled interneurons across the cortical thickness, allowing a fair comparison of input strength across layers. Here, S1 inputs most strongly targeted L5A interneurons. L2/3 SOM+ neurons only received about 30% the input of L5A SOM+ cells (Fig. 5c, p<0.05). L5B was not significantly weaker. L6 SOM+ neurons received the least input (p<0.01). As with PV+ neurons, S1 input was generally perisomatic, especially for L2/3 SOM+ cells. However, the center of mass of the input was shifted below the soma towards white matter for SOM+ cells in L5A, 5B, and 6 (Fig. 5d-f), suggesting that S1 axons were capable of targeting different subcellular domains in different interneuron types.

### PO targeting of PV+ and SOM+ interneurons in M1

Next, we repeated these experiments while labeling thalamic inputs from PO nucleus of thalamus, a higher order somatosensory nucleus whose input to M1 pyramidal neurons had been characterized (Hooks *et al*., 2015; Hooks *et al*., 2013). Thalamic injections were determined in individual animals to be in PO and not anterior motor nuclei by the presence of axonal fluorescence in L1 and L5A. The absence of L5B fluorescence in these animals confirmed that our injections did not target motor thalamic nuclei, such as the anterior components of the ventroanterior-ventrolateral complex, which had previously been shown to arborize in three distinct bands (L1, L5A, and L5B) instead of two (Hooks *et al*., 2013). Targeting recordings to tdTomato+ PV+ interneurons within the arborization of PO axons, we found that, in contrast to S1 inputs, neurons in L5A received the strongest synaptic input (Fig. 6c). However, input to PV+ cells in the top three layers of motor cortex (L2/3, L5A, and L5B) was generally strong, without significant differences. Only synaptic input to L6 was weaker (P<0.05). This input was not centered on the soma in the dendritic arbor, but instead was generally targeted to the deep (white matter) side of proximal dendrites (Fig. 6d-f). This was the case for all four layers measured (L2/3, L5A, L5B, and L6).

PO inputs to SOM+ interneurons were more difficult to characterize. We used the same approach (Fig. 7) to quantify synaptic input. However, the pattern of input to SOM+ was noisy and did not match any previous pattern observed for the other GABAergic cells (Fig 4–6) nor pyramidal neurons previously recorded (Hooks *et al*., 2013; Mao *et al*., 2011). Instead, we found that L5B was the most strongly excited layer (Fig. 7c), with significantly reduced input in L5A (p<0.001). Comparisons to L2/3 and L6 were not significant and showed quite different patterns across different cells, with some pairs favoring L2/3 or L6 and others favoring L5B. The subcellular localization of input was also heterogeneous, with some SOM+ neurons in L5B and L6 receiving input deep to the soma.

### Complementary patterns of input to specific interneuron types

By comparing synaptic strength across layers and normalizing excitability to the strongest input layer for multiple cell types, we were able to compare the pattern of input for the same pathway to multiple cell types. We plotted the normalized input strength graph on the same axes for S1 inputs or PO inputs to PV+ and SOM+ cells, as well as the same data grouped by input to PV+ or SOM+ cells (Fig. 8a-d). This enabled a direct comparison of S1 and PO input to PV+ and SOM+ neurons. Here, the same presynaptic afferents preferred interneurons of different types in different layers. Considering the data organized by postsynaptic target, cortical inputs from S1 to PV+ cells strongly preferred the uppermost (L2/3) PV+ cells, while the functional targeting of PO inputs was biased toward deeper layers (L5A and L5B; Fig. 8c). Input to SOM+ neurons showed a similarly complementary pattern, with input from S1 axons most strongly exciting L5A, which is the layer least strongly excited by PO (Fig. 8d). Thus, the same type of interneuron is targeted in a different manner by the cortical and thalamic inputs. For comparison, we plotted the input strength to excitatory neurons (Hooks, 2017). This pattern shows that S1 and PO generally target pyramidal neurons in the same layer (Fig. 8e). We summarize all the synaptic strength experiments schematically in Fig. 8f, where arrow thicknesses to specific cell types indicate cell-type specific connectivity.

**Figure 8.**
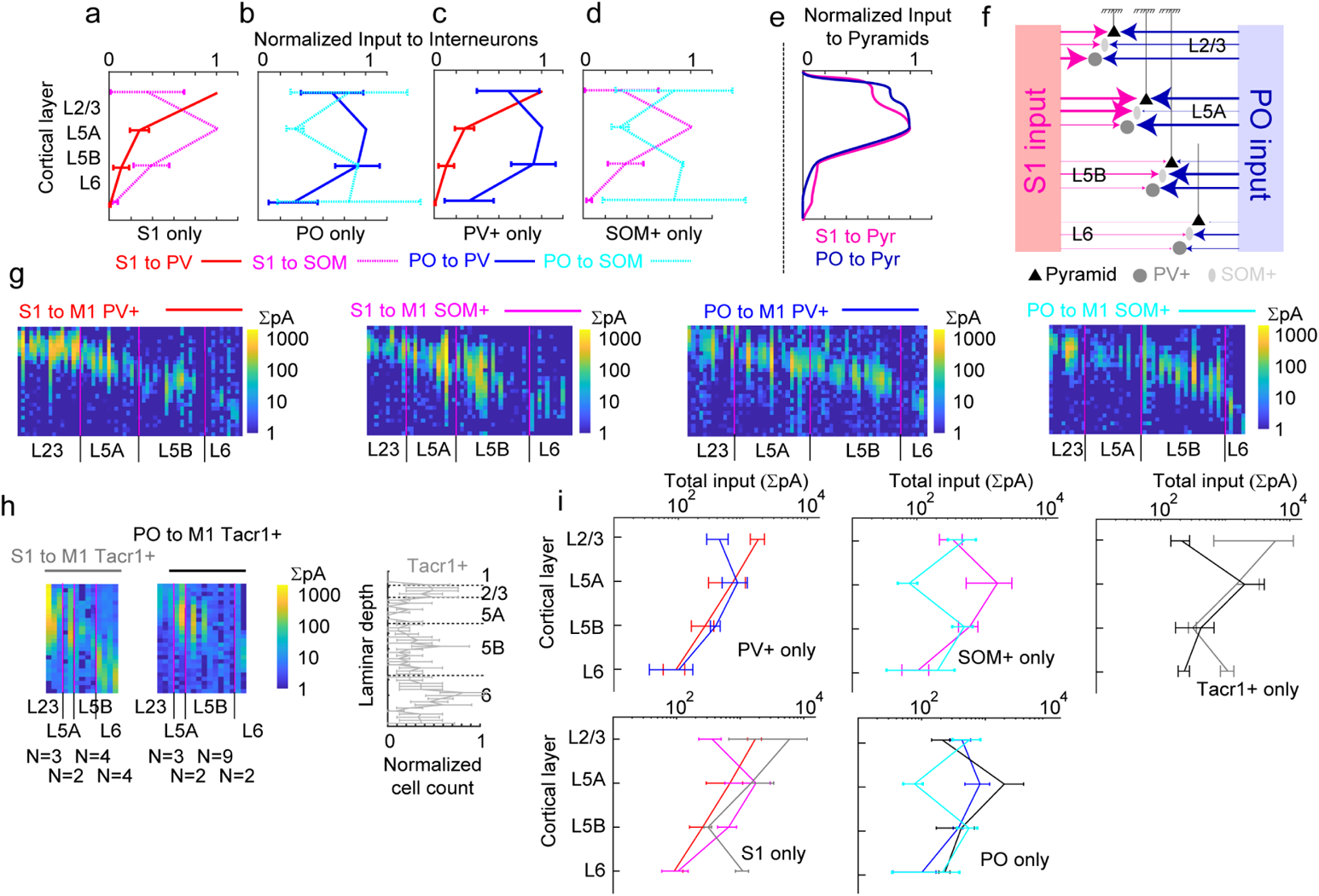
Complementary patterns of input to specific interneuron types. (a-d) Summary graphs comparing the distribution of excitatory input to PV+ and SOM+ neurons. These plots compare normalized input from (a) S1 only or (b) PO only to PV+ and SOM+ neurons. (c-d) compare input to PV+ only (c) or SOM+ only interneurons. (e) The pattern of normalized input strength to pyramidal neurons (Pyr) from S1 and PO, after Mao et al (2011) and Hooks et al. (2013). (f) Cartoon summarizing input maps. (g) Absolute input strength for S1 and PO inputs, plotted as input vectors on a log scale. From left to right, S1 input to PV+ and SOM+ neurons and PO input to PV+ and SOM+ neurons. (h) Tacr1+ neurons (a subset of SOM+ neurons) presented in a similar fashion. Left, input maps from S1 and PO. For Tacr1+ neurons, N=3/2/4/4 for S1 input to L2/3, L5A, L5B, and L6; N=3/2/9/2 for PO input. Normalized laminar distribution of Tacr1+ somata (N=4 slices) measured at right. (i) Absolute input strength averaged by layer and plotted on a log scale for comparison between S1 and PO inputs (top row) to different interneuron types. Comparison of either S! or PO input to different interneuron types presented on the same axes (bottom row).

Because SOM+ neurons comprise a diverse range of neurons, including Martinotti and non-Martinotti subsets (Tremblay *et al*., 2016), we were curious whether the input pattern to the SOM+ neurons labeled in our SOM-Cre x Ai14 strategy would be able to be replicated in a more targeted subset of cells (Huang et al., 2016; Tasic et al., 2018). We labeled the Tacr1+ subset of interneurons using a NK1R-CreER mouse crossed to a tdTomato reporter. Like SOM+ and PV+ crosses, this resulted in labeled neurons across the cortical thickness. We repeated our input mapping experiments in these mice and presented the data in a similar format (Fig. 8h). In contrast to all SOM+ and PV+ neurons, Tacr1+ cells responded well to S1 inputs across multiple layers. Of note, the strongest responses were in in L2/3 and L6, with weaker input to L5 (Fig. 8f-h). The input from PO also did not correspond to the SOM+ pattern, with strongly responding neurons in L5A and L5B.

Because our inputs to PV+ and SOM+ neurons were quantified as normalized input strength, we wondered whether the absolute strength of input to PV+ and SOM+ differed across pathways. We compared input across pathways and cell types by plotting the absolute input strength for all cells on the same scale, as other investigators have done (Kinnischtzke et al., 2014). This was done by summing the EPSC amplitudes across the points of the input map (as in Fig. 3l). Cell vector plots were presented as before (Fig. 8g) using the same logarithmic scale for all plots to improve presentation of more frequent recordings with weaker inputs. In general, excitatory input from S1 and PO to either interneuron type (PV+ or SOM+, top and middle row) was of similar magnitude. Averaging input by layer confirmed that monosynaptic input was of comparable strength (Fig. 8i). The overall pattern is similar to our normalized data. The distribution of S1 and PO input to PV+ is nearly overlapping, with the exception of input to L2/3 PV+ neurons. Input to SOM+ neurons is of similar strength from S1 and PO except for L5A, where the absence of thalamic input is pronounced. Corticocortical input is similar or stronger to SOM+ neurons than PV+ neurons at each laminar depth except L2/3. Thalamocortical input is stronger to L5A PV+ neurons than to SOM+ cells. Outside of this layer, however, input strength to SOM+ cells is equal to PV+ neurons.

### Direct comparison of synaptic and anatomical connectivity

By performing comparable and anatomical experiments, we sought to make a quantitative connection between input to PV+ and SOM+ neurons as measured by these distinct methods. Differences in how the data are quantified presented challenges. First, because the presynaptic neurons labeled by monosynaptic tracing might connect to a starter cell in any cortical layer, we considered our anatomical tracing data to potentially present an averaged view of connectivity to specific interneurons across all cortical layers. To make this data comparable to our physiological data where target PV+ and SOM+ neurons were more explicitly identified by laminar position, we sought to derive an average connection strength to PV+ or SOM+ neurons by converting the layer-by-layer strength of synaptic connectivity (Fig. 9a, left) into a weighted average strength (9a, at bottom) using the fractional distribution of PV+ and SOM+ neurons (potential starter neurons) in each layer (Fig. 9a, middle) as weighting factors. This not only determined the weighted average, but also by resampling the individual neurons recorded in each category, a bootstrap mean (±SEM) could be determined to assess variability of this estimate of physiological connectivity. The weighted averaged gave an idea of the relative input strength to PV/SOM from cortical (0.7842) and thalamic (1.3724) input. This suggested that monosynaptic thalamic input favored PV+ neurons, while corticocortical input might favor SOM+ neurons, likely because the strongest recipient layers for SOM+ (L5A and L5B) contained larger numbers of cells that the strongest PV+ recipient layer (L2/3). A ratio of PV/SOM input based on anatomical tracing (Fig. 9b) could also be determined. We quantified the mean fraction of presynaptic input in somatosensory cortex or thalamus. Using this as an anatomical connection strength, we then made a ratio between PV and SOM as done for synaptic input. Anatomical input from both somatosensory cortex (PV/SOM, 1.2045) and thalamus (1.0853) slightly favored PV+ neurons in M1. To explore the degree to which these ratios varied, we compared N=1000 bootstrap replicates of the synaptic and anatomical data, finding largely overlapping distributions (Fig. 9c).

**Figure 9.**
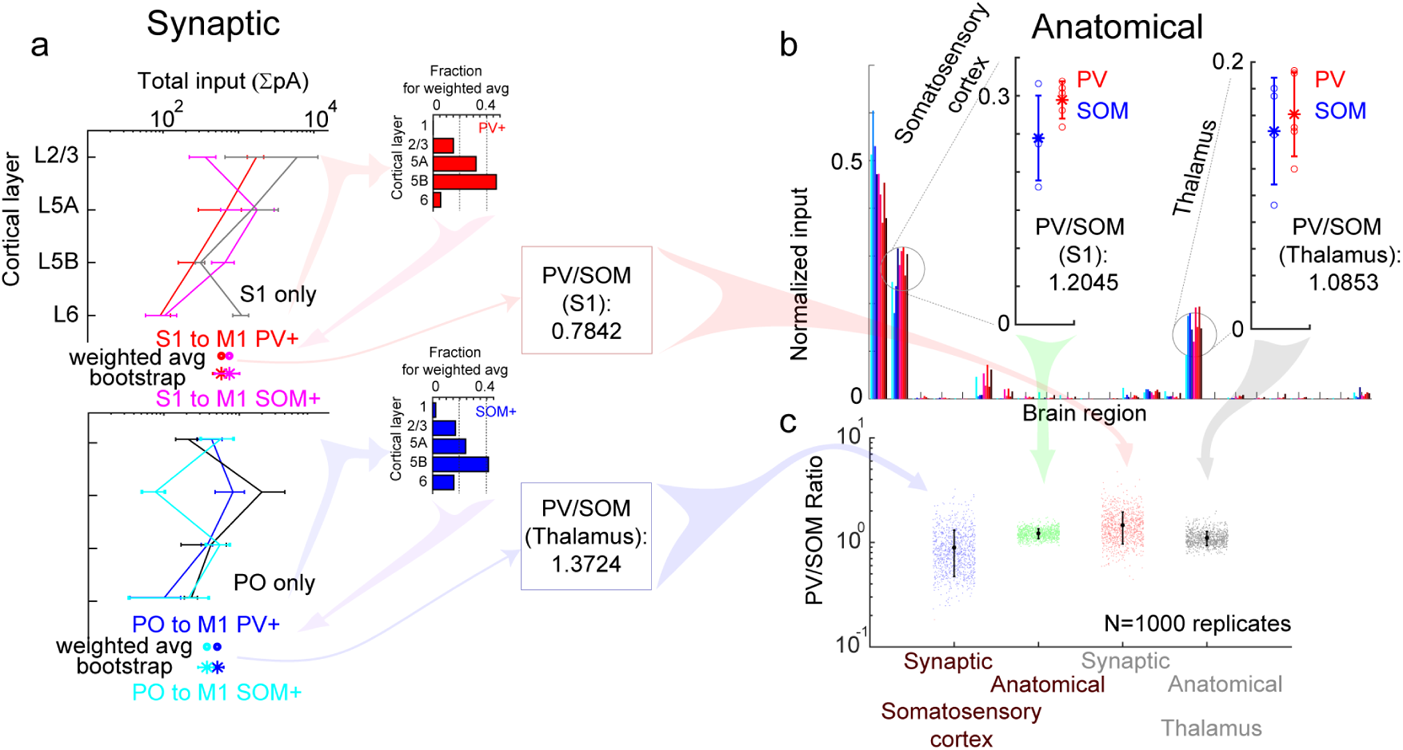
Direct comparison of synaptic and anatomical connectivity. (a) Left, absolute synaptic input strength plotted for comparison between S1 inputs to PV+ and SOM+ neurons. Right, the distribution of interneurons per layer. The weighted average input strength is shown below the graph, either as a single point (weighted avg) or the mean and SEM from N=10000 replicate (bootstrap) analysis. The ratio of input to PV/SOM is shown for each plot. (b) Comparison of anatomical input to PV+ (N=5) and SOM+ (N=4) neurons in M1 from retrograde tracing (after Fig. 1). Insets show normalized fraction of input from each case. The ratio of input to PV/SOM is shown for each plot. (c) Comparison of PV/SOM input for synaptic inputs (from S1, red; from PO, green) and for anatomical tracing (in S1, blue; in thalamus, gray). The data was resampled for N=1000 replicates to compare.

### Short term plasticity of long-range excitation to PV+ and SOM+ neurons

Our measurements of synaptic strength were based on monosynaptic input under recording conditions (TTX, CPP, 4-AP) which monitor synaptic responses to the first stimulus. These measurements might not reflect the relative contribution of the two interneuronal populations to a train of stimuli, which might be more physiological. Since PV+ and SOM+ neurons show differences in short term plasticity of local excitatory inputs (Beierlein et al., 2003), we were interested to test whether later synaptic inputs in a train might alter which interneuron population was more likely to carry feedforward inhibition.

In contrast to these results in PV+ neurons, SOM+ cells showed some degree of facilitation for both classes of input. S1 input was more facilitating at 20 Hz than 10 Hz. Similarly, PO input was also more facilitating at 20 and 40 Hz. These experiments were performed using the same viral vector and recording conditions for both mouse lines, suggesting that the short-term plasticity observed differed between postsynaptic targets and was not due exclusively to an artifact of measuring short term plasticity with optical methods (Jackman et al., 2014). We do not know whether the same axon might have different release properties depending on its cellular target (Beierlein *et al*., 2003; Gabernet et al., 2005), or whether different long-range axons from these areas preferentially innervate PV+ and SOM+ neurons.

## DISCUSSION

Anatomical models for cortical connectivity propose that potential functional connectivity is proportional to the overlap of axons and dendrites (Braitenberg and Schuz, 1998; Peters and Feldman, 1976). However, a range of excitatory and inhibitory neurons exist in cortex (Tasic *et al*., 2018; Tremblay *et al*., 2016), each with potentially cell type-specific rules for connectivity and plasticity. Here we show that cortical and thalamic long-range inputs to M1 excite two types of inhibitory interneurons in different and complementary ways. S1 inputs most strongly targeted L2/3 PV+ neurons, but SOM+ neurons in middle layers (L5). Further, while S1 and PO both targeted pyramidal neurons in L2/3 and L5A with similar connection strength, these pathways targeted PV+ neurons in a different fashion. These results indicate that cortical and thalamic afferents form highly specific connections in neocortex. Thus, the recruitment of feedforward inhibition may not be simply a blanket of inhibition, but instead is divided into specific feedforward circuits.

What is the functional relevance of specificity in feedforward connectivity? If incoming afferents activate a selected subset of interneurons, then certain inputs will activate some cortical networks while silencing others. For M1, this might mean activating the appropriate response to incoming sensory stimuli, while inhibiting others. This would be of use, for example, in selection or inhibition of one action over others. Pyramidal neurons in different cortical layers project to different (but partially overlapping) long range targets, including corticocortical, corticostriatal, corticothalamic, and pyramidal tract type projections (Harris and Shepherd, 2015; Network, 2021). Thus, we hypothesize differences in laminar inhibition can shape the pattern of M1 output to subsets of these targets. For example, S1 activation of L2/3 PV+ neurons might be expected to most strongly silence output of L2/3 pyramids, resulting in reduced feedforward excitation in both local inputs to L5 pyramidal neurons (Hooks *et al*., 2011; Weiler *et al*., 2008) as well as long-range corticocortical connections (Harris and Shepherd, 2015). This would vary during longer trains of stimuli as synapses to PV+ neurons depress. Notably, while both S1 and PO pathways excite L5B more weakly than other layers (Hooks *et al*., 2013; Mao *et al*., 2011), PO excited both PV+ and SOM+ interneurons in this layer, which might aid in terminating signals associated with inappropriate motor responses. Similarly, short-term plasticity differences in facilitation and depression may shape the timing and duration of inhibition, including low-pass or high-pass filtering of feedforward signals (Blackman et al., 2013). Thus, M1 computations performed on feedforward excitatory inputs from S1 and PO may be shaped by the relative amounts of facilitating and depressing inhibition recruited.

### Specific circuits for feedforward inhibition

Motor cortex integrates a wide range of input from frontal cortical areas (Dum and Strick, 2002; Reep *et al*., 1990; Rouiller *et al*., 1993), primary and secondary sensory cortex (Chakraibarti and Alloway, 2006; Hoffer *et al*., 2003; Kaneko *et al*., 1994a; Smith and Alloway, 2013; Suter and Shepherd, 2015), retrosplenial cortex (Yamawaki et al., 2016), and different thalamic nuclei (Deschenes *et al*., 1998; Kuramoto *et al*., 2009; Ohno *et al*., 2012; Strick and Sterling, 1974). These inputs activate M1 during voluntary and sensory guided movement, with each targeting excitatory neurons in M1 in a layer-specific manner (Hooks *et al*., 2013). Different M1 neurons can represent either sensory input or motor behavior such as whisking, licking, or lever pressing (Harrison and Murphy, 2014; Huber *et al*., 2012; Masamizu et al., 2014; Tanaka et al., 2018), suggesting that individual neurons are parts of different networks, receiving input of different strength from cortical inputs. Here we tested whether this pattern is the same for functional synaptic input to interneurons, finding that the laminar pattern of excitation varies with the postsynaptic cell type targeted.

Each excitatory pathway evokes feedforward inhibition to some degree (Isaacson and Scanziani, 2011). This disynaptic inhibition might serve to terminate continuing excitation from the same inputs. Once recruited, local circuit interneurons connect to large numbers of nearby pyramidal neurons (Bock et al., 2011; Fino and Yuste, 2011; Packer and Yuste, 2011). If synaptic weights are strong, this might provide an effective suppression of cortical activity (Fino et al., 2013), though the effectiveness of these connections could be reweighted by activity-dependent (Xue et al., 2014) or experience-dependent (Chen et al., 2015) plasticity mechanisms. Thus, it is of interest to understand whether the recruitment of feedforward inhibition proceeds through distinct interneurons (of either the same or different type) for inputs recruiting motor cortex. Our data indicates that there is space for selectivity in inhibitory circuitry.

Some specific examples of differences stand out in our quantitative maps. Our data suggests that the incoming sensory information from S1 excites PV+ neurons in L2/3 but SOM+ neurons in L5. Since these same afferents from S1 also strongly target pyramidal neurons in L2/3 and L5A (Mao *et al*., 2011), it suggests that, in part, the feedforward inhibition recruited to L2/3 pyramids and L5A pyramids can proceed through different neural populations, as cortical interneurons generally, though not exclusively, inhibit pyramidal neurons in the same layer (Katzel *et al*., 2011). Since SOM+ neurons may also inhibit PV+ cells, helping to disinhibit cortex (Pfeffer et al., 2013), it is possible that S1 input can have differential inputs on the excitability of L2/3 and L5A pyramids, despite similarity in monosynaptic excitation to these targets. This suggests that differences in cell types play a role in determining the efficacy of feedforward input and the rules of long-term plasticity in these two target layers. The pattern of thalamic input to PV+ interneurons also differed from S1 input. This suggests that where cortical and thalamic afferents target the same excitatory network, to the extent S1 and PO target partially non-overlapping PV+ populations, feedforward inhibition recruited by one pathway would not be as strongly depressed by the recruitment of PV+ inhibition by the other pathway arriving at a similar time.

Of note, PO provided fairly strong input to both PV+ and SOM+ neurons in L5B (Fig. 8g-h). This feedforward inhibition would potentially inhibit L5B pyramidal neurons, which are only weakly excited by S1 and PO L5A (Hooks *et al*., 2013; Mao *et al*., 2011). We infer the net effect would be to reduce activity in L5B pyramidal neurons. It is unknown if there is specific targeting to certain pyramidal neuron subtypes. L5B contains two main classes of output neurons, IntraTelecephalic type cells projecting to both cortex and striatum, as well as Pyramidal Tract type cells which mainly target thalamus, brainstem, and spinal cord (Hooks et al., 2018; Shepherd, 2013). Thus, a range of cortical output channels might be suppressed. The function of inhibiting neurons outside the normal targets of excitation of these pathways is unknown, but by reducing overall activity of M1 output neurons might serve to set a higher threshold of activation in the context of incoming sensory information. By reducing background activation, this would increase the signal to noise ratio of outgoing motor signals.

### Organization of long-range input to SOM+ interneurons differs across cortical areas

It had previously been shown that SOM+ neurons in L4 of sensory areas generally received weaker input than neighboring pyramidal neurons and PV+ neurons (Bruno and Simons, 2002; Cruikshank *et al*., 2007; Cruikshank *et al*., 2010). This thalamocortical preference for PV+ over SOM+ neurons in somatosensory cortex extends to L2/3 (Naskar et al., 2021). Thalamic input to SOM+ does exist in sensory areas, though it may be developmentally downregulated at ages younger than those studied here (Marques-Smith et al., 2016; Tuncdemir et al., 2016). Thus, we were surprised to find many layers in M1 where SOM+ cells received comparable inputs to PV+ cells (Fig. 8) as measured by monosynaptic input mapping (Porter et al., 2001). Comparisons are challenging due to limitations of normalizing input strength across different injections in different animals (Petreanu *et al*., 2009), but the mean thalamic input strength to SOM+ interneurons was marginally higher than that to PV+ neurons except for L5A (Fig. 8h middle right). However, there is considerable variability between individual cells (Fig. 8g). Similarly, corticocortical input from S1 was higher to SOM+ neurons in L5A and L5B (Fig. 8h top right). This collectively suggests that there is no uniform rule that thalamic input will be selective for PV+ neurons across all cortical areas and layers.

In addition to monosynaptic input strength, it is also worth considering that input from S1 and PO will not occur in isolation but as part of trains of action potentials, especially should thalamus fire in burst mode (Sherman, 2001; Steriade et al., 1993). In the case of PO, short term depression of input to PV+ neurons is the strongest short term plasticity we observed (Fig. 10). In contrast, these inputs were facilitating to SOM+ neurons. This suggests that as rapidly as the second or third pulse in a train, the recruitment of feedforward inhibition could shift from PV+ to SOM+ neurons (Tan et al., 2008). The difference in short term facilitation and depression to SOM+ and PV+ neurons respectively is not as large for S1 afferents as for PO inputs, though the general pattern is the same. This effect is qualified by differences in the intrinsic excitability of the particular cells involved, as well as their own depression onto pyramidal neuron targets. Inhibition from SOM+ neurons is hypothesized to be facilitating, in contrast to the depression observed in fast spiking neurons (Beierlein *et al*., 2003). If this is the case in M1, then this would serve to further shift the balance of disynaptic feedforward inhibition from PV+ to SOM+ neurons, giving SOM+ neurons the potential to play a role in regulating excitability and plasticity (Chen *et al*., 2015).

**Figure 10.**
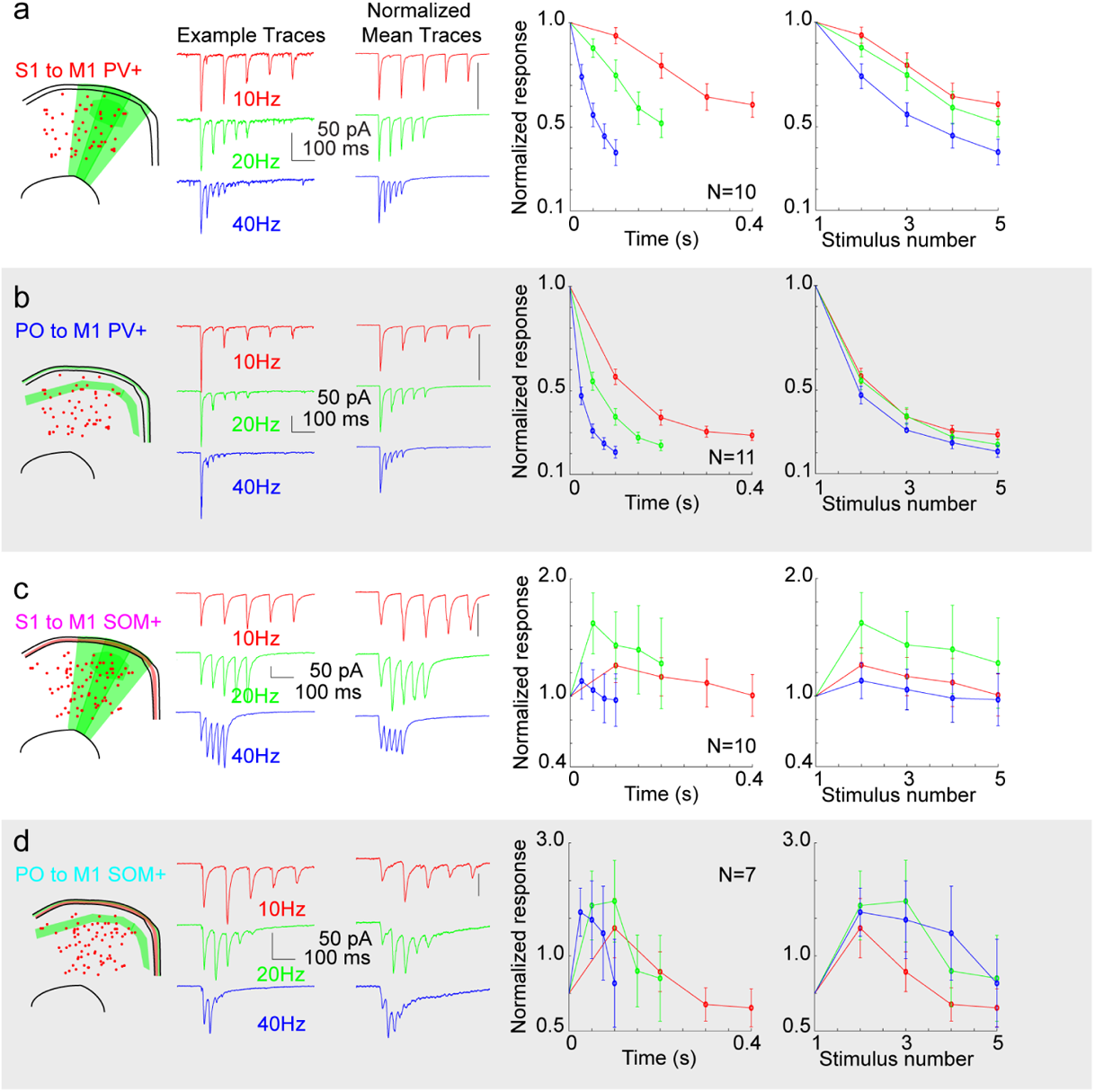
Short-term plasticity at long range inputs to interneurons of mouse motor cortex. (a) Short term plasticity of S1 input to PV+ neurons. Left, example traces showing PV+ neuron response to S1 stimulation at 10 Hz (red), 20 Hz (green), and 40 Hz (blue) stimulation of Chronos-GFP+ S1 axons. Scale, 50 pA and 100 ms. Middle, average normalized traces for the same stimulus frequencies. Scale bar=1.0. Right, summary graphs showing normalized responses for 10 Hz, 20 Hz, and 40 Hz. Graphs are plotted based on time of the pulses (right) or the stimulus number (far right). (b) Short term plasticity of PO input to PV+ neurons, plotted as in (a). (c) Short term plasticity of S1 input to SOM+ neurons, plotted as in (a). (d) Short term plasticity of PO input to SOM+ neurons, plotted as in (a).

### Comparison of anatomical and functional methods for mapping microcircuitry

The emergence of techniques for recording, imaging, and reconstructing the whole brain present a great deal of data to deepen our understanding of how neocortical areas are connected and function at large scales. Imaging projections from defined neurons (Jeong et al., 2016; Oh *et al*., 2014; Zingg *et al*., 2014) to label long-range targets or retrograde viral methods to label presynaptic inputs (Luo et al., 2018; Wickersham et al., 2007a) offers hope for efficiently identifying long-range connections in whole brains when images are aligned to standardized coordinate systems (Eastwood et al., 2019; Hooks *et al*., 2018; Kuan *et al*., 2015). Thus, monosynaptic tracing with rabies virus has substantial appeal. Ideally, a comparison across large scale recording methods (such as probes with numerous channels or imaging calcium responses in large neuron populations), synaptic connectivity, and anatomical projections would help clarify how neural connectivity contributes to brain function.

Here we directly compared synaptic input and retrograde anatomical tracing of presynaptic input to M1 interneurons. Based on prior work, we hypothesized that thalamocortical input would favor PV+ interneurons. In S1, thalamic inputs to fast spiking (presumed PV+) neurons were more likely to be connected (Bruno and Simons, 2002) and were vastly stronger by EPSC amplitude, as much as 10-20x in L4 and L5/6 (Fig. 6, (Cruikshank *et al*., 2010)) as well as later 2/3 (Fig. 2, (Naskar *et al*., 2021)). In M1 by contrast, both synaptic connectivity mapping experiments and retrograde viral tracing data showed strong thalamocortical connections to PV+ and SOM+ neurons in M1 (Figs. 1 and 6-9). Indeed, input to PV+ neurons was only ~10-30% stronger than that to SOM+ neurons (Fig. 9). This difference was not significant in the anatomy data, though non-significant trends (Fig. 1) were present in three thalamic nuclei not assessed functionally (VM, PC, and PF). Thus, both measures of anatomical and functional connectivity were consistent in demonstrating distinct thalamocortical connectivity rules for primary motor cortex compared to sensory cortex. Similarly, our measures of corticocortical input to SOM+ neurons were comparable between anatomical and functional methods (Fig. 9), though the functional method showed a slight preference for SOM+ neurons (because of stronger input to the more numerous interneuron population in L5A and L5B), while the anatomical method showed a nonsignificant trend for larger S1 input to PV+ neurons (Fig. 1). This is in contrast to corticocortical input in S1, where long range input to PV+ neurons was stronger in L2/3 for multiple pathways (Naskar *et al*., 2021).

However, there are some limitations to the comparison. With anatomical methods, it was not possible to label SOM+ or PV+ interneurons in specific layers and show differences in connectivity as was the case in functional mapping experiments. Furthermore, quantifying differences in synaptic strength is not precisely the same as labeled presynaptic neurons. Although we hypothesize that stronger synaptic connections would result in more points of contact between corticocortical or thalamocortical axons and rabies-infected starter cells, there are not control experiments to show that increased synaptic strength results in an increase in labeled presynaptic neurons. Furthermore, while we can quantify synaptic strength to PV+ or SOM+ neurons from S1 or PO inputs in a given layer, it is not certain that deriving a single value for connection strength for the whole population is the strongest means to make such a comparison.

How reliable is retrograde labeling for anatomically assessing connectivity? The data presented here suggest anatomical and functional methods can predict similar connectivity. First, the retrograde tracing data provides strong support for retrograde tracing as an effective measure of area-to-area connectivity. Tracing from starter cells in each cortical area specifically labeled cortical areas and a pattern of thalamic nuclei unique to that cortical area and reproducibly across replicates (Fig. 2b-e). For example, M2 gets input from limbic areas while M1 and S1 do not. Furthermore, even small brain areas, such as individual thalamic nuclei, have distinct patterns of label for each of the three cortical areas studies. However, because the specific thalamic labeling was almost identical for input to PV+ and SOM+ interneurons for the regions that we studied, evidence that retrograde tracing varies strongly with postsynaptic target would be better to test in cell types with vastly different connectivity, especially where afferents avoid innervating one cell type while connecting strongly to another intermingled type.

### Subcellular organization of input to interneurons

Our circuit mapping approach gives some insight into the subcellular localization of synaptic input in the dendritic arbor of postsynaptic neurons. This insight is constrained by methodological limitations of patch clamp which limit our ability to study distal dendrites. Interneurons, however, are generally more electronically compact than pyramidal neurons. There is some organization of the input around the PV+ and SOM+ soma, suggesting some mechanisms for afferent targeting exist and may differ between interneuron subtypes. S1 input to PV+ neurons is evenly distributed perisomatically, while these same afferents tend to target dendrites ~50-150 μm below the soma of SOM+ cells. Similarly, PO input to PV+ cells is also shifted deep to the soma center by ~50-150 μm. This offset is interesting in contrast to thalamic inputs to pyramidal neurons (Petreanu *et al*., 2009). In these cells, PO input targets the apical dendrites, and especially the L1 arbors of L3 pyramidal neurons but is soma-centered (and quite strong) for L5A pyramidal neurons. This is consistent with each cell type directing input from a defined presynaptic pathway to some extent within its arbor.

### Different SOM+ subtypes have different connectivity

One difficulty in making a general statement about SOM+ connectivity in M1 was the variability within this dataset, especially for thalamic inputs (Fig. 7). In our assessment of monosynaptic input strength and short-term plasticity, PV+ neurons behaved in a much more homogeneous manner. These are the largest GABAergic neuron population defined by a single immunohistochemical marker (~40%), consisting of mostly perisomatic-targeting basket cells, though this group also includes axon-targeting chandelier cells (Tremblay *et al*., 2016). In contrast, SOM+ neurons (~30%) include distal dendrite targeting Martinotti-type neurons, evident in the L1 axonal fluorescence in Fig 3f, as well as multiple non-Martinotti types. Thus, SOM+ neurons include at least two (and likely more) molecular subtypes (Ma et al., 2006; Naka et al., 2019; Tasic *et al*., 2018; Zeisel et al., 2015), which might account for the variability in the data. One subset of SOM+ interneurons expresses Tacr1, as well as NPY and nNOS (Kubota et al., 2011), though these have differences in nNOS expression level (Dittrich et al., 2012). Tacr1+ cells, like both PV+ and SOM+ neurons are present across cortical layers 2-6. This made comparison of input strength possible. This was not the case for VIP+ neurons, which are much more numerous in only upper layers (~60% in L2/3 (Pronneke et al., 2015)), as well as other non-VIP expressing 5HT3aR+ interneurons which are similarly biased to L1 and L2/3 (Lee *et al*., 2010).

When synaptic strength was mapped, this presented a pattern of input strength quite in contrast to that of the SOM-Cre x Ai14 mice (Fig. 8h-i). In particular, S1 input had been distributed across L2/3, L5A, and L5B to SOM+ neurons, but most strongly to L5A. In contrast, this subset had a quite distinct pattern, with substantial L2/3 and L6 input. PO input was noisy to SOM+ neurons in general. This emphasized the likelihood that each molecular subtype of interneuron had distinct mechanisms to govern synaptic strength.

## Conclusion

Overall, these results point towards circuit mechanisms by which different pathways of incoming excitation may excite specific subcircuits, including specific networks for inhibition in primary motor cortex. Having some means by which recruitment of feedforward inhibition from each pathway does not generally silence all M1 circuits is perhaps necessary for control of movement when M1 is receiving competing inputs from different brain regions.

## MATERIALS AND METHODS

### Stereotactic injections

Animal protocols were approved by Institutional Animal Care and Use Committee at University of Pittsburgh (Protocol #20118160) and followed the recommendations in the Guide for the Care and Use of Laboratory Animals of the National Institutes of Health. Mice of either sex were used. Experimental procedures were similar to previous studies (Hooks *et al*., 2013). Cre driver mouse lines (PV-Cre, SOM-Cre, and NK1R-CreER) were used in conjunction with the lsl-tdTomato reporter line, Ai14, to label specific interneuron populations (see Table 2). Tacr1 experiments using NK1R-CreER, required tamoxifen injections to drive recombination. Animals were anesthetized using isoflurane and placed in a custom stereotactic apparatus. Juvenile mice at postnatal (P) day P12-P22 were injected with AAV expressing excitatory opsins. AAVs included CAG-ChR2-mVenus and hSyn-Chronos-GFP. AAV-CaMKIIa-hChR2(H134R)-EYFP was used in Tacr1 mapping experiments (see Table 3 for details and serotypes). Injections were made with glass pipettes (Drummond) using a custom-made positive displacement system (Narashige). Stereotactic coordinates are listed in Table 1. A pair of injections (50-100 nL; 500 and 800 μm depth from pia) was made in the cortex. One location (50-100 nL) was injected into thalamic targets.

**Table 2.**
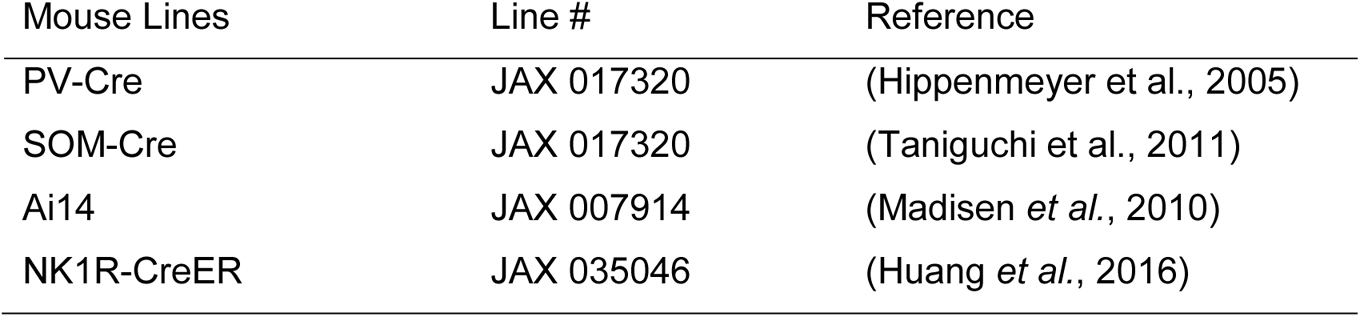
Mouse lines.

**Table 3.** Viral reagents

| Virus | Serotype | Addgene # |
| --- | --- | --- |
| CAG-ChR2-mVenus | AAV1, AAV9, and AAV10 | 20071 |
| hSyn-Chronos-GFP | AAV1, AAV5, and AAV9 | 59170 |
| CaMKIIa-hChR2(H134R)-EYFP | AAV9 | 26969 |
| DIO-TVA-EGFP-B19G | AAV1 | 52473 |
| N2c-ΔG-EnvA-tdTomato | CVS-N2c rabies | n/a |

### Monosynaptic viral tracing

We used AAV injections to express DIO-TVA-EGFP-B19G in potential starter cells in cerebral cortex of PV-Cre and SOM-Cre driver mouse lines (Atasoy et al., 2008; Wickersham *et al*., 2007b). After two weeks of expression time, we injected EnvA pseudotyped rabies (CVS-N2c^ΔG^ strain) into cortex and allowed 8-9 days for transport and expression (Reardon *et al*., 2016). Following transcardial perfusion, whole brains were sectioned, immunoamplified for EGFP and TdTomato, and stained with NeuroTrace Blue (ThermoFisher) as a structural marker. The brain was imaged and reconstructed using NeuroInfo software (MBF Bioscience) as previously described (Eastwood *et al*., 2019; Hooks *et al*., 2018). Original images are posted at: http://gerfenc.biolucida.net/link?l=Jl1tV7 (Biolucida Cloud Viewer, free download) and http://gerfenc.biolucida.net/images/?page=images&selectionType=collection&selectionId=32 (web-based). Images were aligned to the Allen CCF V3.0 (Kuan *et al*., 2015; Oh *et al*., 2014) using NeuroInfo software (MBF Bioscience). Somata were detected by NeuroInfo using an artificial neural network. Their coordinates in the Allen CCF used to quantify in which structure the neuron was detected using custom Matlab software. Presynaptic labeled (tdTomato+) neurons in each structure were quantified as a fraction of total labeled neurons in that brain to normalized for differences between mice and injections. For comparisons between starter cell types (PV+ or Cre+) or injection sites (M2, fM1, vS1), t-tests were performed. To reduce the number of comparisons to a manageable size, brain regions were only included in the analysis if the fraction of presynaptic neurons labeled exceeded a threshold value: 2% for 24 large brain areas (Fig. 1o and Fig. 2bd) or 0.5% for 33 smaller thalamic nuclei (Fig. 1p and 2c and 2e-h).

The two-stage procedure of Benjamini, Krieger, & Yekutieli (Benjamini *et al*., 2009) was used to control for the false discovery rate (FDR). Black * (with p-values noted) mark significant differences between presynaptic labeling. Gray * used for comparisons not significant after correcting for FDR.

### Electrophysiology and photostimulation

Brain slices were prepared >14 days after viral infection, in young adult mice (~P30-P60). Mice were anesthetized with isoflurane and the brain was rapidly removed and placed in cooled choline-based cutting solution (in mM: 110 choline chloride, 3.1 sodium pyruvate, 11.6 sodium ascorbate, 25 NaHCO_3_, 25 D-glucose, 7 MgCl_2_, 2.5 KCl, 1.25 NaH_2_PO_4_, 0.5 CaCl_2_). Off-coronal sections (300 µm) of M1 were cut using a vibratome (VT1200S, Leica). Additional sections were cut to confirm injection location. Slices were incubated at 37 °C in oxygenated ACSF (in mM: 127 NaCl, 25 NaHCO_3_, 25 D-glucose, 2.5 KCl, 2 CaCl_2_, 1 MgCl_2,_ 1.25 NaH_2_PO_4_,) for >30 min and maintained at room temperature (22 °C) thereafter.

Whole cell recordings were performed at 22 °C in oxygenated ACSF with borosilicate pipettes (3-6 MΩ; Warner Instruments) containing potassium gluconate-based internal solution (in mM: 128 potassium gluconate, 4 MgCl_2_, 10 HEPES, 1 EGTA, 4 Na_2_ATP, 0.4 Na_2_GTP, 10 sodium phosphocreatine, 3 sodium L-ascorbate; pH 7.27; 287 mOsm). These recordings targeted neurons in whisker M1 for whiskers (vM1, coordinates in Table 1). In some experiments, biocytin was added to the intracellular solution (3 mg/mL biocytin or neurobiotin). For subcellular ChR2-Assisted Circuit Mapping (sCRACM) experiments (Petreanu *et al*., 2009), TTX (1 µM, Tocris), 4-AP (0.1-0.3 mM, Sigma) and CPP (5 μM, Tocris) were added to the bath. Dose-response of 4-AP concentration was confirmed in a subset of experiments comparing EPSC amplitude while 4-AP was added. Under these conditions, laser pulses (1 ms) depolarized ChR2-expressing axons in the vicinity of the laser beam and triggered the local release of glutamate. Measurements of postsynaptic currents then revealed the presence of functional synapses between ChR2-expressing axons and the recorded neuron near the photostimulus. Blocking action potentials prevented polysynaptic contributions.

Data were acquired at 10 kHz using an Axopatch 700B (Molecular Devices, Sunnyvale, CA) and Ephus software (www.ephus.org) (Suter et al., 2010) on a custom-built laser scanning photostimulation microscope (Shepherd et al., 2003). During sCRACM mapping, neurons were held at −70 mV. A blue laser (473 nm, CrystaLaser) was controlled via scan mirrors (Cambridge Technology). Light pulses were controlled with a shutter (4 ms open time) in series with an acousto-optic modulator (1 ms pulse, Quanta-Tech) to deliver ~0.5-2 mW at the specimen plane through a low power objective (4x, 0.16 NA, UPlanApo; Olympus). Laser power was constant during a given day. Sweep consisted of 100 ms baseline and 300 ms following onset of the stimulus. The map grid (12×26 sites at 50 μm spacing) was centered horizontally over the soma of the recorded neuron, aligned at its upper edge to the pia, and covered the entire interneuron dendritic arbor. Maps were repeated 2-4 times and averaged across trials. Stimuli for short term plasticity were delivered using 1ms flashes of a 470 nm LED (Cairn). Sweeps were interleaved and repeated 4-6 times per stimulus frequency.

### Data analysis

Data analysis was performed with custom routines written in Matlab. Electrophysiology data were low pass filtered (1 kHz). EPSCs were detected with a threshold of >6x standard deviation from baseline). Mean EPSC amplitude for input maps was averaged over 75 ms post-stimulus (reported in pA). For comparison between cells in the same slice, total input was computed by summing supratheshold pixels. Paired comparisons across cells use the nonparametric Wilcoxon signed-rank test. Mean ratio of excitation were calculated as geometric means, with the strongest input layer shown as 1.0. In these comparisons, the bar graph was overlaid with a line showing the bootstrap mean and SD (10000 replicates) resampling the paired data. To compare the axonal profile of S1 and PO axons (Fig. 1), we quantified the fluorescence as before (Hooks *et al*., 2013) by determining the mean fluorescence of pixels at a given laminar depth (binned in 50 bins across cortex from pia to white matter). N are reported in figures for number of cells. Number of mice used are: N=15 (S1 to M1 PV+), N=12 (PO to M1 PV+), N=10 (S1 to M1 SOM+), N=10 (PO to M1 SOM+), N=4 (S1 to M1 Tacr1+), and N=4 (PO to M1 Tacr1+).

---

**Supplemental Figure 1.**
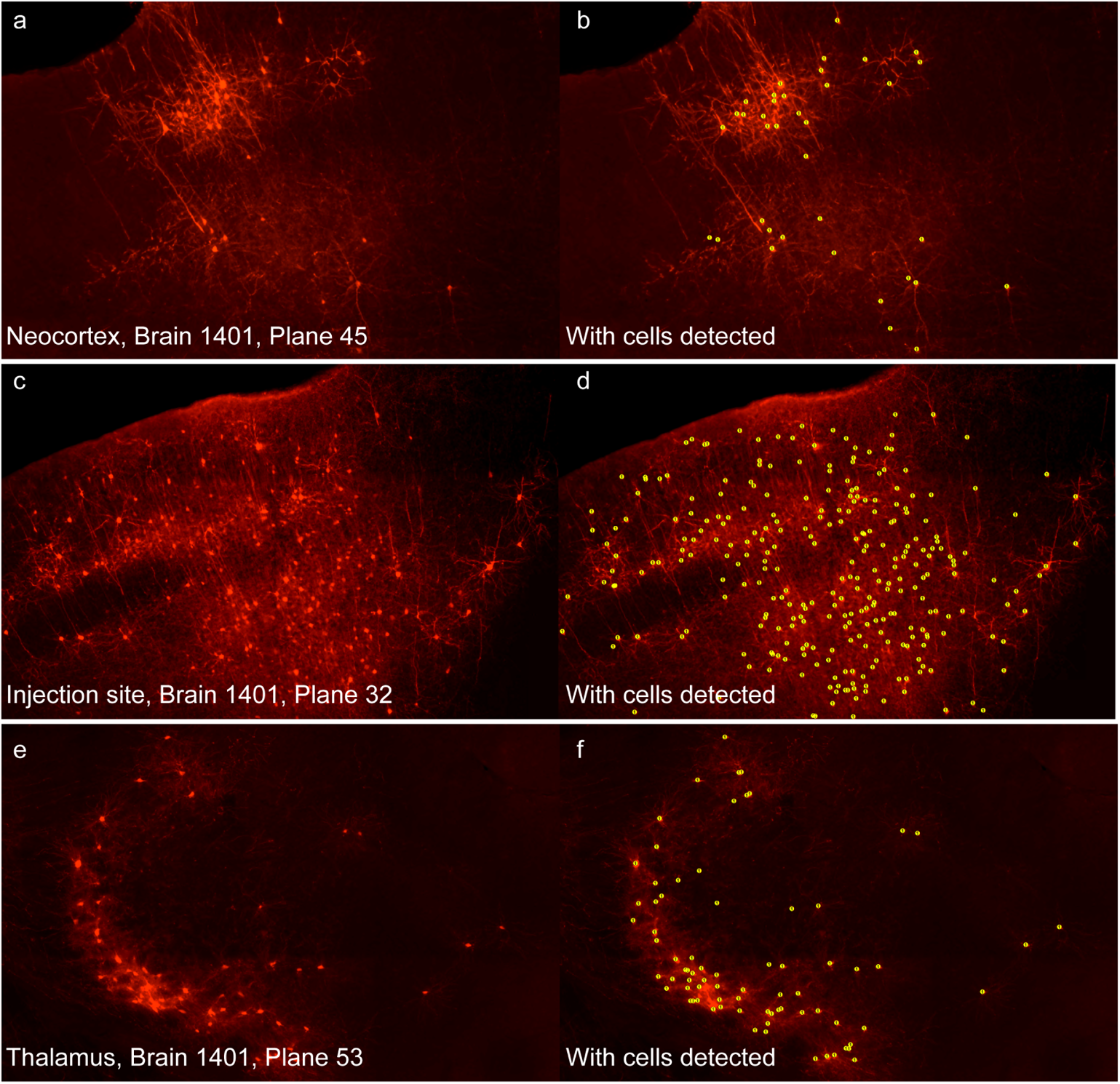
Cell detection in cortex and thalamus. (a-b) Low density cortical labeling from monosynaptic rabies tracing in M1 of a SOM-Cre mouse. (a) Cells in somatosensory neocortex posterior to the injection site. (b) The same plane with cells detected. (c-d) High density cortical labeling near the injection site in M1 of a SOM-Cre mouse. (e-f) Thalamic labeling in a SOM-Cre mouse.

**Supplemental Figure 2.**
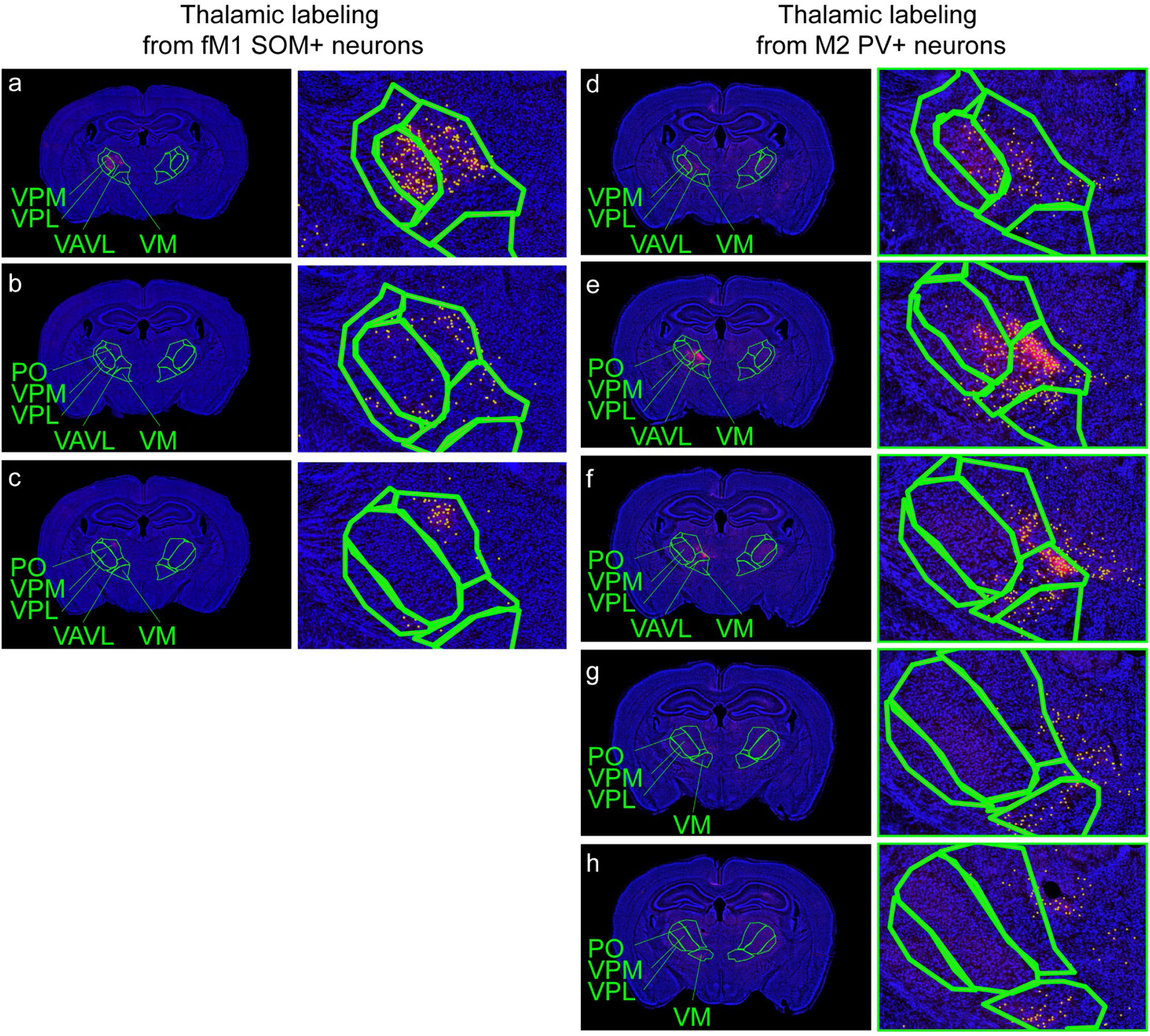
Thalamic labeling in from SOM+ and PV+ neurons. (a-c) Retrograde labeling from fM1 SOM+ neurons. Left panel, structural stain (Neurotrace Blue) and retrogradely labeled neurons (tdTomato, red). Thalamic nuclear boundaries based on registration to CCF V3.0 atlas. Nuclei as indicated. Right panel, higher magnification version. Detected cells labeled by yellow circles. Planes shown are separated by ~0.16 mm proceeding from anterior (top, a) to posterior (bottom, c). (d-h) Retrograde labeling from M2 PV+ neurons. Left panel, structural stain (Neurotrace Blue) and retrogradely labeled neurons (tdTomato, red). Thalamic nuclear boundaries based on registration to CCF V3.0 atlas. Nuclei as indicated. Right panel, higher magnification version. Detected cells labeled by yellow circles. Planes shown are separated by ~0.16 mm proceeding from anterior (top, d) to posterior (bottom, h). PO, posterior thalamic nuc.; VAVL, ventoanterior/ventrolateral thalamic nuc.; VM, ventromedial thalamic nuc.; VPL, ventroposteriolateral thalamic nuc.; VPM, ventroposteriomedial thalamic nuc.

**Supplemental Figure 3.**
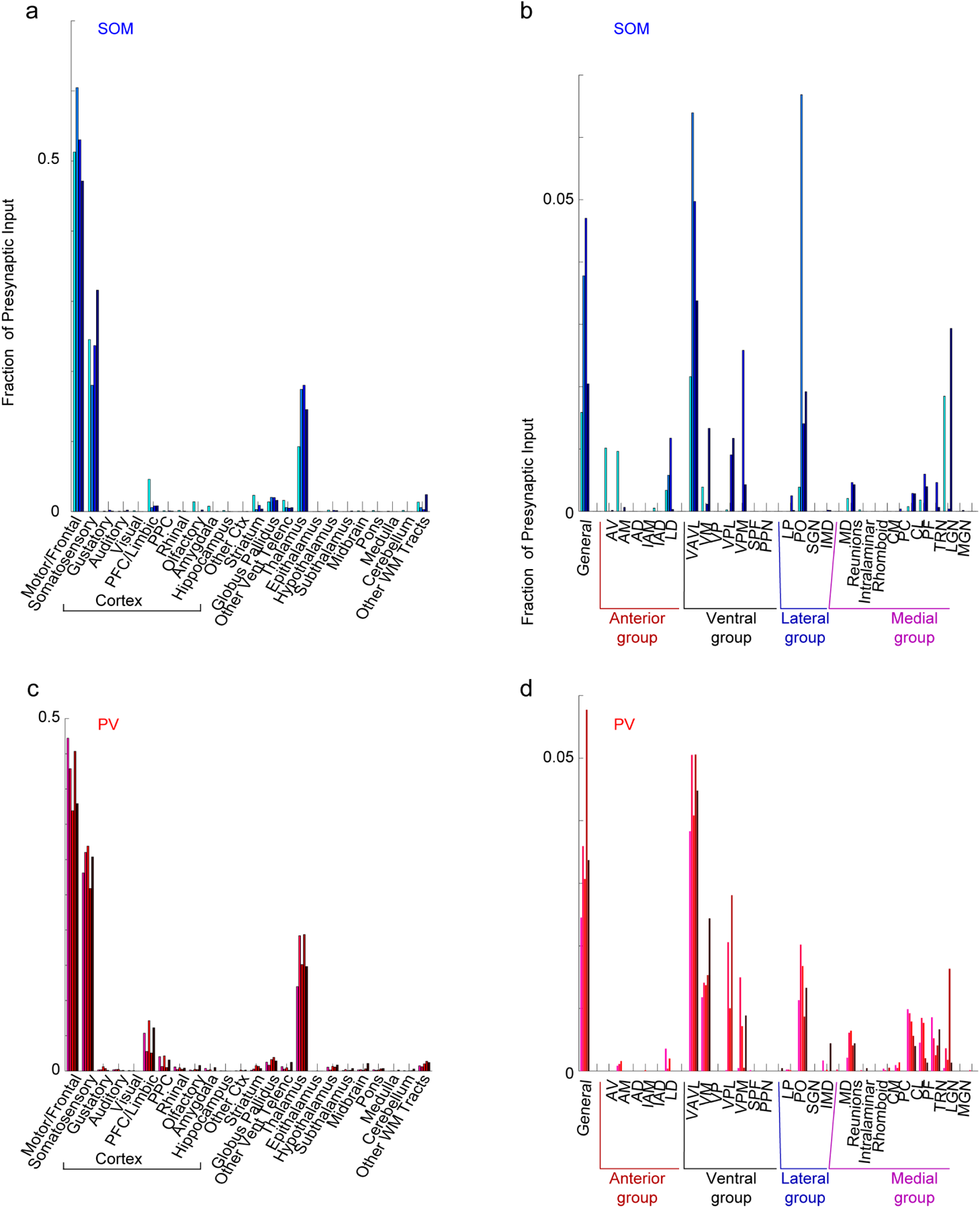
Consistency of monosynaptic retrograde tracing. (a-b) Quantification of presynaptic neuron location in the whole brain (a) and thalamus (b) plotted as bar graphs to emphasize differences between individual retrograde tracing cases. For SOM-Cre (N=4, blue) mice, the fraction of the total presynaptic neuron population (±SEM) plotted for all M1 injections. Bars represent individual cases. (c-d) Quantification of presynaptic neuron location in the whole brain (a) and thalamus (b) for PV-Cre (N=5, rec) mice, plotted as above.

**Supplemental Figure 4.**
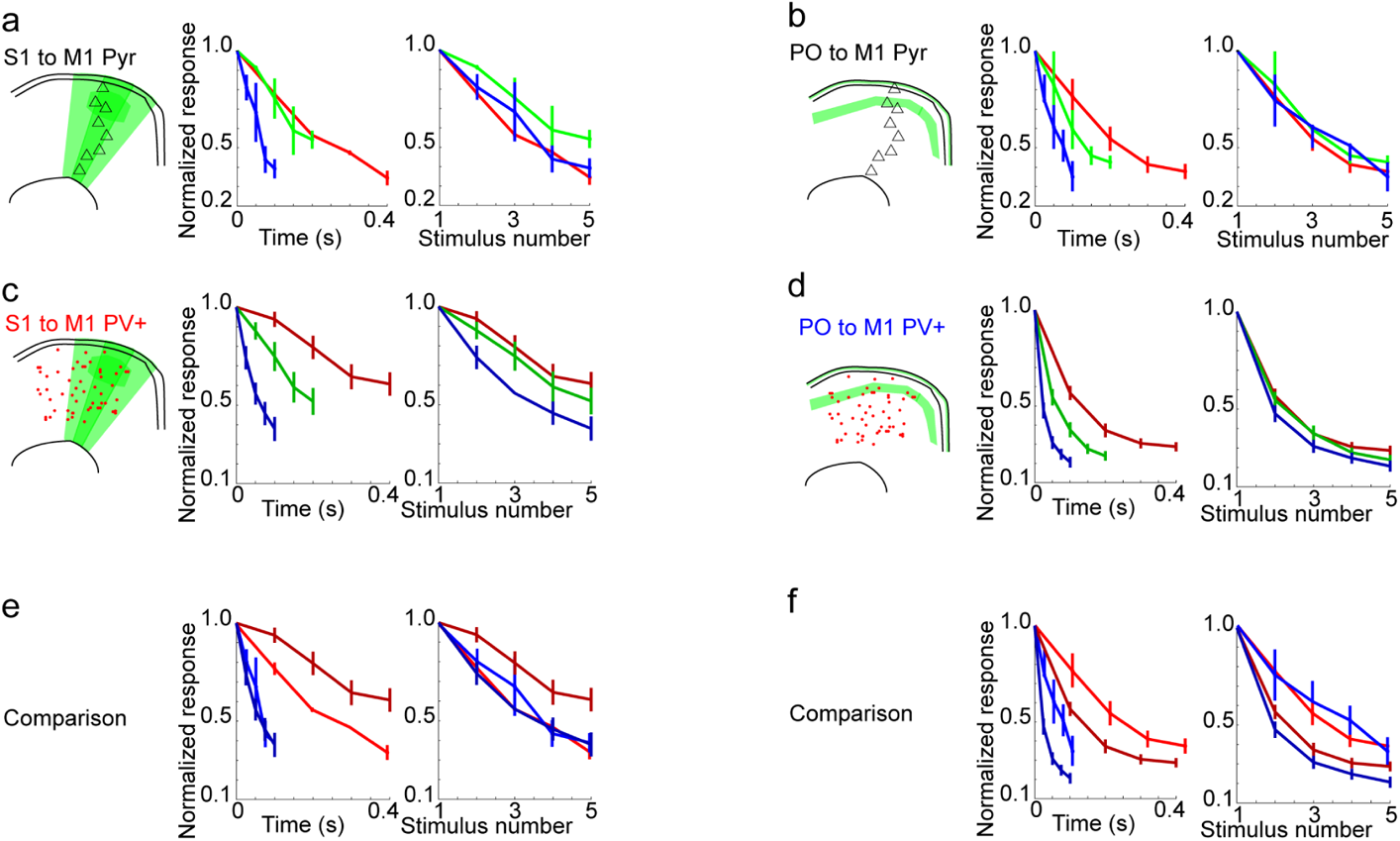
Short-term plasticity comparisons of PV+ interneurons and pyramidal neurons. (a) Short term plasticity of S1 input to pyramidal neurons. Left, normalized response to S1 stimulation at 10 Hz (red), 20 Hz (green), and 40 Hz (blue) stimulation of Chronos-GFP+. Summary graphs show responses plotted based on time of the pulses (left) or the stimulus number (right). (b) Short term plasticity of PO input to pyramidal neurons, plotted as in (a). (c,d) Short term plasticity of S1 input (c) and PO input (d) to PV+ interneurons plotted for comparison. (e,f) Comparison plot of the above data for 10Hz and 40Hz stimulation on the same axes.

## ACKNOWLEDGEMENTS

We thank Caroline Runyan, Chinfei Chen, Nuo Li, Srivatsun Sadagopan, Taehyeon Kim, Shelby Ruiz, and other members of the Hooks lab for comments and suggestions. Jay Couey performed whole cell recordings contributing to the circuit mapping and short-term plasticity experiments. This work was supported by a NARSAD Young Investigator Award (BMH), NIH NINDS R01 NS103993 (BMH), and a CDMRP PRMRP Discovery Award PR201842 (RG and BMH). The authors declare no competing financial interests or other conflict of interest.

